# DNA-barcode labelled MHCII multimers for detection of antigen-specific CD4 T cells across large libraries of epitopes

**DOI:** 10.64898/2026.06.23.733927

**Authors:** Yogesh Basavaraju, Suzan Dijkstra, Tripti Tamhane, Signe Koggersbøl Skadborg, Liying Lu, William W Kwok, Lawrence Stern, Georg M. Lauer, Sine Reker Hadrup

## Abstract

The role of antigen-specific T cells responding to antigen is a topic of intense studies, and critical for mechanistic insight of diseases and development of therapeutic strategies. Methods for broad-scale detection of antigen-specific CD4 T cells are lacking, while such methods have demonstrated great value in exploring CD8 T cell response in health and disease. Furthermore, major histocompatibility complex II (MHCII) assays are technically challenging due to high HLA diversity, lower binding affinities, low frequencies of *ex vivo* antigen-specific CD4 T cells and several bottlenecks in production and peptide exchange of MHCII monomers.

Here we use peptide-loaded MHCII (pMHCII) proteins multimerized on a barcode- and fluorophore-labelled dextran backbone to provide a method for the detection of peptide-specific CD4 T cells by using a large display of MHCII-associated peptides. We have established a protocol for MHCII production and peptide-exchange suitable for the generation of large libraries of peptide-MHCII complexes. We validate the use of such pMHCII complexes in the form of barcode-labelled MHCII multimers to detect antigen-specific CD4 T cells. We demonstrate that we can identify antigen specific CD4 T cells, using these DNA barcoded peptide-MHCII multimer. The multimer bound CD4 T cells were selected based on the fluorochrome signal, and the co-attached DNA barcodes were hereafter amplified and used to identify the peptide-MHCII response/binding. In cases where the peptide-specific CD4 T cells frequencies are very low, we expanded the cell population with peptide-pools and in the presence of IL2. The given CD4 T cell populations hereby reach a cell number allowing for the DNA-barcoded pMHCII multimers to detect responses otherwise missed out.

Applying this technology, we utilized a panel of 150 peptides derived from human cytomegalo virus (CMV), Epstein barr virus (EBV), Influenza (Flu), SARS CoV 2 and SARS CoV1, Hepatitis B virus (HBV), and Hepatitis C virus (HCV) loaded onto HLA-DRB1*01:01 and DRB1*04:01 to screen peripheral blood mononuclear cells (PBMC). We assessed *ex vivo* responses in 16 participants with HCV infection, and successfully detected naturally occurring viral-specific CD4 T cells at frequencies as low as 0.004% of total CD4 T cells. The low-frequency responses, identified via the barcode screen, were rigorously validated using individual fluorophore-labelled tetramer staining after a peptide-driven expansion in 15 participants. Furthermore, we assessed the recognition of novel HCV epitopes in 11 additional participants. Through this, we identified a total of 12 distinct HCV epitopes, including 9 that have not been previously utilized in assays to detect CD4 T cells.

Overall, this barcoded-multimer platform provides a powerful tool for the large-scale discovery of class II epitopes and the broad profiling of CD4 T cell specificities. This method will allow for in-depth analyses of immune interactions, provide a better understanding of the antigen-driven associations between CD4 and CD8 T cell responses, and help dissect the complexities of CD4 T cell protection in HCV infection.

## Introduction

T cells are one of the main drivers of adaptive immunity, and have been studied and exploited for therapeutic interventions in infectious diseases, cancer, and autoimmunity ^1^. T cells are activated when their T cell receptors (TCR) recognize their cognate antigen in the form of a peptide presented on the respective major-histocompatibility complex (MHC) molecule along with co-stimulatory signals from the presenting antigen-presenting cell (APC) and their surroundings ^2^. Detecting such antigen-specific T cells is important for the design and optimization of therapies and vaccinations. During the last decade, substantial development has lifted the technologies available for the detection of CD8 T cells, from low-throughput ^3^, model antigen approaches, to large peptide-MHC library screens, allowing for the assessment of CD8 T cell recognition of large antigen pools, such as viruses ^4 5^ and cancer antigens ^6^. CD4 T cells are key mediators of adaptive immunity and are an important factor in orchestrating the immune response. CD4 T cells are known to assist by B cell activation and antibody maturation along with helping to activate cytotoxic CD8 T cells ^7^. CD4 T cells have also been found to exhibit cytotoxic activity in cancer patients 14,15,^10^. Therefore, identification of antigen-specific CD4 T cells is important in many immune-related diseases and therapies. However, the technologies developed to assess CD4 T cell responses ^11 12^ have not yet been adapted to enable the identification of peptide-specific CD4 T cells on a large scale. Consequently, most CD4 T cell analyses rely on functional evaluation, i.e. activation, proliferation, including cytokine secretion in response to antigens ^13^. While such assays can be of great value, they require large amount of PBMC material and are not easy adaptable to screening for T-cell recognition against large antigen libraries ^14 15^. To allow such screening for peptide-specific CD4 T cells, direct assessment of their peptide-MHCII (pMHCII) binding capacity is desirable ^16^. Click or tap here to enter text.Click or tap here to enter text.Click or tap here to enter text.Click or tap here to enter text.

HCV represents a major global health challenge, with an estimated 50 million people living with chronic HCV infection worldwide, resulting in approximately 240,000 deaths annually from complications such as liver cirrhosis and hepatocellular carcinoma ^17^. While acute infection is spontaneously cleared by ∼25% of individuals, the remaining ∼75% progress to chronic disease^18,19^. Though direct-acting antiviral therapies with high cure rates are available, there is no prophylactic vaccine and an estimated 1 million new infections still occur annually ^17^. The development of an effective vaccine is hindered by the virus’s high genetic diversity and an incomplete understanding of the immune function within the context of HCV^20 21^. CD4 T cells are crucial for viral clearance, where both resolvers and chronically infected patients mount comparable CD4 T cell responses, and it is the functional quality rather than the quantity that determines the infection outcome^22^. As the current patient screening heavily relies on low-throughput functional assays or fluorophore-tetramer assays with the limited library of class II HCV epitopes that is available, there are significant gaps in the ability to map the landscape of MHC class II HCV epitopes required to inform vaccine design.

Soluble MHCII expression and purification have long been challenging. However, recent advances has enabled the successful production of a broad range of MHC class II alleles ^23^. Recombinant expression of MHCII started in E.coli as inclusion bodies ^24^, then insect cells were used ^25^, but both these methods had low yields of the product or could not be applied across many MHCII alleles. Currently, MHCII is expressed with a covalently bound CLIP peptide in mammalian cells with the possibility of intracellular biotinylation ^26^. The soluble MHCIIs can hereafter be purified, multimerized and used to detect CD4 T cell responses, based on fluorescent labeling, which by combinatorial coding can increase the number of responses that are evaluated ^16^. Improvement in strategies to exchange the MHCII covalently bound to CLIP with a peptide in the presence of HLA-DM, has allowed for generation of large libraries of pMHCII variants ^27^. Thus, with the innovation in MHCII expression, purification and peptide exchange strategies, we now have the possibility to develop large-scale screening approaches to determine the peptide-MHCII recognition of CD4 T cells.

Click or tap here to enter text.Click or tap here to enter text.Click or tap here to enter text.Click or tap here to enter text.Here we present a high-throughput method to screen for antigens and detect antigen-specific CD4 T cells in PBMC as both direct *ex vivo* and peptide expanded. Fluorescence based markers limit the scalability of methods for CD4 T cell interrogation, therefore we used DNA barcodes as identification tool for peptide-MHCII multimers and identify CD4 T cell binding. The method was tested for specificty, repeatability, sensitivity, and detected responses were validated with conventional tetramers stains. Thereafter, we applied the DNA-barcoded pMHCII multimer technology to screen HCV patient samples (n=16) with upto 150 viral peptides from other known viruses and HCV and detected new epitopes of relevance for CD4 T cell recognition of HCV. In total, we found 12 HCV peptide-responses, out of which 9 are novel. These novel epitopes were validated with conventional tetramer stains and could be extended to patients outside the cohort. Thus, the barcoded-pMHCII multimer platform is a tool for large-scale discovery of MHCII epitopes across virus and other diseases. The epitopes identified could pave the way for vaccine testing in virus infections, detecting CD4 T cell epitopes in tumor-infiltrating lymphocytes (TILs), and aid in understanding the interplay between CD8 and CD4 T cells in health and disease.

## Results

### Antigen-specific CD4 T cells can be captured and identified using barcode-labelled pMHCII multimers

DNA-barcoded peptide-MHCII (pMHCII) multimer method of identifying antigen-specific CD4 T cells begins with the expression of soluble DRB1*01:01-CLIP in mammalian cells, purified, biotinylated, and then peptide-exchanged with different peptides binding to the MHCII allele to create a stable pMHCII complex. This pMHCII complex is then mixed with barcode- and fluorophore-labelled, streptavidin-conjugated dextran backbone to generate a pMHCII multimer. Each peptide-specific multimer is represented by a unique barcode in a pool of multimers used to stain the cell mixtures, after which the pMHCII multimer positive CD4 T cells were sorted based on the fluorophore conjugated to the dextran backbone, and the pMHCII-associated DNA barcodes were amplified by PCR, sequenced, and analyzed using Barracoda 2.0 (**Figure 1A**). Peptide-MHCII dependent, barcode-enriched responses are plotted using a log fold change (logFC) value, with specific T cell response detection defined as a logFC of greater than 2 and a p value of <0.001.

**Figure 1:**
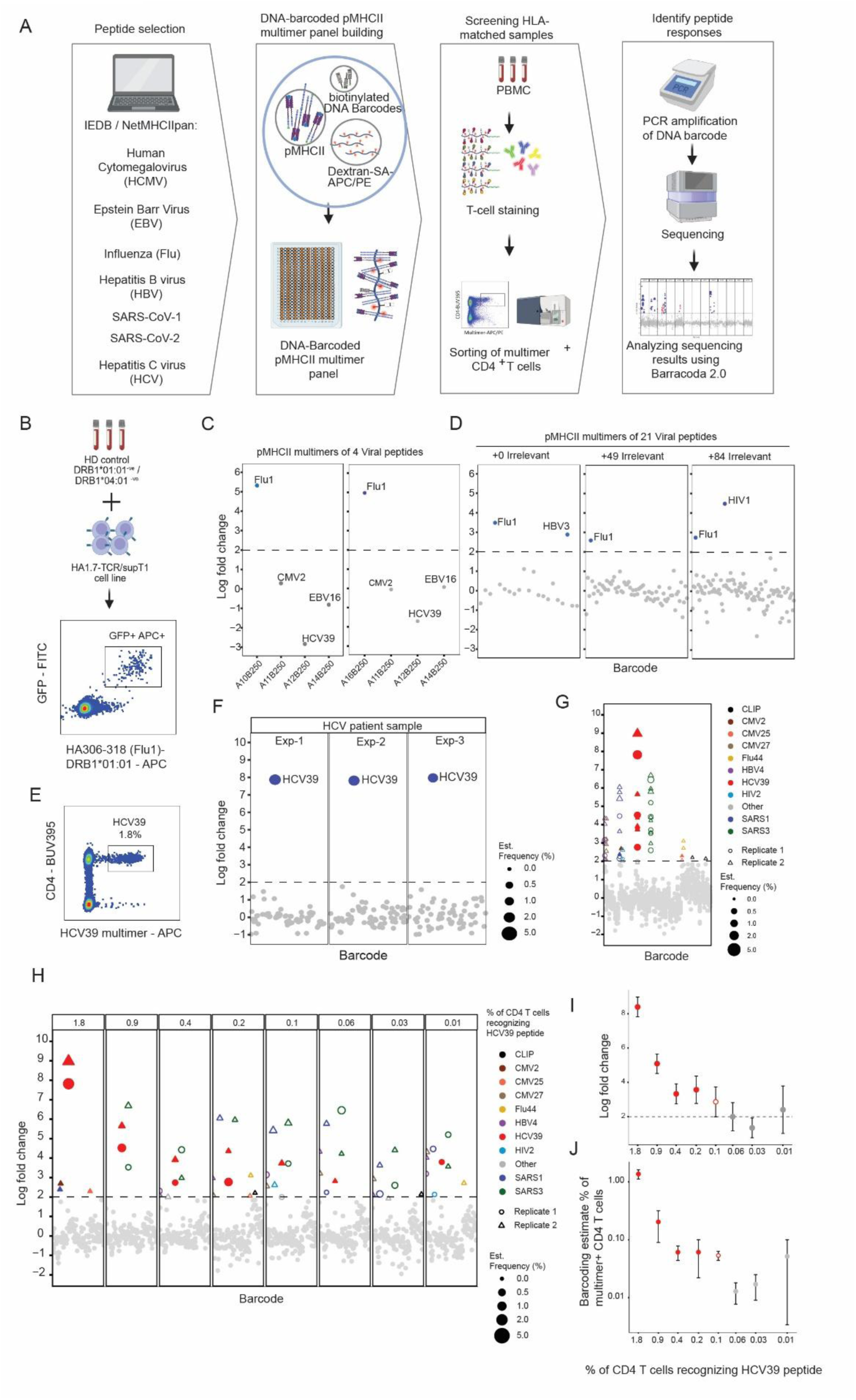
Methodology and testing of DNA-barcoded pMHCII multimers-. A) Schematic overview of the methodology using DNA-barcoded pMHCII multimers to detect antigen-specific CD4 T cells. B) HA1.7-TCR engineered supT1 cells (HA1.7-TCR/supT1, GFP+), having affinity towards HA_306-318_(Flu1) peptide, were spiked into PBMC from a DRB1*01:01 negative donor. Testing of Flu1-DRB1*01:01 (DRB1) multimer by detecting the spiked HA1.7-TCR/supT1 cells which are GFP^+^ and multimer positive. C) Detection of HA1.7-TCR/supT1 cells using pMHCII-barcode multimers generated with 4 viral peptides, each denoted by their respective barcodes. Two different sets of barcoded multimer panels were used (left and right side, respectively). The blue dot marks the detection of significant barcode-enriched signal, in this case Flu1. The Y axis shows the log-fold change (logFC) depicting the fold-change in barcode amplification for all -pMHCII within the library. All pMHCII with barcode enrichment, logFC>2 (represented by dotted line) and p<0.001, are marked as significant T cell responses detected in the given sample towards the given pMHCII. D) Detection of spiked-in HA-TCR/supT1 cell responses based on three barcoded multimer panels containing 21, 70, and 105 pMHCII multimers, respectively, each comprising 21 virus-derived pMHCII including Flu1 peptide; and 0, 49, and 84 irrelevant pMHCII (DRB1-CLIP) respectively with each one of the pMHCII having unique barcodes. E) Flow plot of HCV39^+ve^ CD4 T cells detected by the DRB1*01:01-HCV39 dextan multimer in HCV patient 1. F) Detection of HCV39 specific CD4 T-cell responses in PBMC 08-023 using a panel of 70 barcoded pMHCII multimers. Data is displayed for 3 individual experiments. G) Enrichment of DNA barcodes within the sorted multimer positive CD4 T-cell population, following analyses of samples using 70 barcoded-pMHCII multimers (21 viral peptides + 49 irrelevant peptides). The plot shows responses in 8 samples. The samples are a 2-fold dilution series of HCV patient 1 PBMC diluted in HD donor PBMC (not-HLA typed). Each dot represents a unique pMHCII-associated barcode. The dotted line represents significance threshold for the barcode enrichment (logFC>2 and p<0.001). The peptides are colored based on the virus that they are derived from and the size of the dots represent the estimated frequency from the analyses. The replicates are shown in circles and triangles for replicate 1 and 2, respectively. Significant responses detected in donor Patient 1 are marked as filled dots, and other responses (from non-HLA typed donor) are marked as outline. H) Each individual dilution of PBMC 08-023 is displayed, with the theoretical frequency of HCV39 specific T cells written on top of each plot, while the estimated frequency from barcode analysis is depicted as the size of the point. The replicates are shown as circles and triangles for replicate 1 and 2, respectively. I) Correlation between the theoretical frequency (x-axis) and log fold change (logFC) of the barcode enrichment (y-axis) for the HCV39 response in Figure H. The dotted line represents the significance level. The dot is filled with red when the logFC of both replicates is greater than 2; outlined red when the logFC of only one replicate is greater than 2 and the other is close to 2 (>1.90); and colored grey when one or both of the logFC of the duplicates is less than 1.90. J) Correlation between the theoretical frequency (x-axis) and the estimated frequency based on barcode enrichment and frequency of multimer^+^ CD4 T cells (y-axis). The dots are colored as per the color scheme in Figure 1I.

We first wanted to test the ability of a single pMHCII multimer to detect its respective CD4 T cells in a pool of multimers. We generated a control cell line by engineering parent supT1 cells to express a TCR (HA1.7) specific to the HA_306-318_ (Flu1) epitope, which is presented in the context of DRB1*01:01 and DRB1*04:01 (**Supplementary figure 1**). The engineered supT1 cells (HA1.7-TCR/supT1), which also have green fluorescent protein marker (GFP+), were added to PBMC from a DRB1*01:01 and DRB1*04:01 negative healthy donor and subsequently tested with Flu1-multimer (**Figure 1B**). We then generated 5 pMHCII multimers from peptides including CMV2, EBV16, and HCV39, and 2 representing Flu1; all with unique barcodes. The 5 pMHCII multimers were then made into two sets of four, each containing CMV2, EBV16, and HCV39 multimers, and one of the Flu1 multimer. The two sets of multimers were then used to screen HA1.7-TCR/supT1 cells spiked into a DRB1*01:01 and DRB1*04:01 negative donor. We successfully identified Flu1-specific barcodes being enriched and shown as significant responses (logFC > 2, p<0.001). No other pMHCII multimer was enriched, thereby establishing the use of pooled barcode-labelled pMHCII multimers to specifically detect a given CD4 T cell response (**Figure 1C**). We hereafter wanted to explore the detection of the Flu1 specific population using larger pMHCII multimer panel. To do this we build 3 panels comprising 21, 70 and 105 peptides derived from common viral proteins including Flu1. Each of the panels consisted of 21 viral peptide (**Supplementary table 1**) and the rest were CLIP-associated MHCII (DRB1*01:01), each of them represented by a unique barcode. Each of the 3 panels were then used to screen HA1.7-TCR/supT1 cells spiked into a DRB1*01:01 and DRB1*04:01 negative donor. HA1.7-TCR/supT1 cells were consistently detected in all three settings, demonstrating that the library size did not compromise peptide-specific barcode enrichment (**Figure 1D**). We also observed significant enrichment of barcodes associated to other viral pMHCII, HBV3 and HIV1, that may represent unspecific binding of the pMHCII complexes in question. Yet, the rest of the peptides including pMHCII-CLIP as irrelevant peptide controls were not detected, suggesting the method does work efficiently to sort out the Flu1 response. We would get a better representation of the method if we applied it with endogenous CD4 T cell detection, rather than using a cell line.

Having established the functionality of the assay, we next sought to test the method’s ability to detect endogenous CD4 T cells. We used PBMC from a participant with HCV infection and a known CD4 T cell response against epitope HCV39 (1.8% of total CD4 T cells, Figure 1E). Using a panel of 70 barcoded-labelled pMHCII multimers against Flu, CMV, EBV, SARS CoV including HCV39, we screened the HCV patient sample, across three independent experiments. The HCV39-specific barcodes were enriched consistently and the HCV39 response was detected in all three replicates with a similar logFC value. We then checked the barcode-estimated frequency, which is an estimated frequency of peptide-specific CD4 T cells out of the multimer-positive CD4 T cells sorted. The estimated barcode frequency of HCV39-specific CD4 T cell response was 1.9%, 1.5%, and 2.2%; which closely aligns to the actual frequency of 1.8% of total CD4 T cells (**Figure 1F**).

To establish a sensitivity range we made a dilution series of the HCV39 response detected in sample 08-023. The original frequency of 1.8% HCV-specific CD4 T cells was diluted 2 fold in a non-HLA typed PMBC. Eight samples were generated with a theoretical range of 1.8% to 0.01% HCV39-specific CD4 T cells out of total CD4 T cells. These samples were then evaluated in duplicates for detection of HCV39-specific CD4 T cells using the barcode-labelled pMHCII multimer panel of 21 viral peptides and 49 irrelevant peptides. We detected barcode enrichment specific to HCV39 pMHCII, along with other peptides derived from Flu, CMV, and SARS-CoV (**Figure 1G**). The original PBMC 08-023 showed 3 additional responses, i.e. SARS1, CMV2, and CMV25 peptides, suggesting these could be additional viral CD4 T cell responses not previously tested for in this sample. In the sample dilutions, in addition to the expected HCV39-specific CD4 T-cell responses, we observed CD4 T cell recognition towards four other peptides including SARS3, HBV4, and CMV27 in the majority of the samples (**Figure 1H**). Three of these responses were exclusively detected upon diluting the PBMC 08-023, suggesting these responses are derived from the PBMC used for dilution. Decreasing the HCV39-specific CD4 T cell frequency by dilution effectively decreased the logFC of the enriched barcodes representing the HCV39 peptide. At 0.1% of theoretical HCV39-specific T cell frequency, we see one of the duplicates enriched, and the other close to the threshold (logFC = 1.99) suggesting 0.1% could be the lower limit of detecting antigen-specific CD4 T cells using the DNA barcoded-pMHCII multimer method (**Figure 1I**). Based on these experiments we conclude that pMHCII-multimer binding CD4 T cells can be detected in PBMC samples when the frequency is >0.1%, and that frequencies estimated based on barcode analysis correlated well with theoretical frequency for HCV39-specific T cell responses of >0.1% (**Figure 1J**).

### Pre-cultivation with antigenic peptide pool can enhance detection capacity while using barcoded MHCII multimers

To further expand and optimize the use of barcoded-pMHCII multimers in identifying CD4 T-cell populations within previously untested PBMC material, we applied a panel of 70 barcode-labelled pMHCII multimers (comprising 21 viral and 49 irrelevant peptides) to screen for T-cell recognition in PBMC from an DRB1*01:01 positive healthy donor, HD1. The PBMCs were evaluated under three distinct conditions: 1) directly *ex vivo*, 2) peptide-pool expanded with IL2, where cells were expanded with a 21-viral-peptide pool (matching the viral peptides from the 70 barcode-labelled pMHCII multimer panel), and 3) irrelevant expansion, where the same expansion method was used with an irrelevant peptide (HIV1-ACQGVGGPGHKARVLA^28^) instead of the 21-viral-peptide pool (**Figure 2A**). Analysis of the pMHCII-associated barcodes within the sorted T-cell populations revealed significant barcode enrichment corresponding to a number of CD4 T cell responses towards multiple viral peptides in all three conditions. While some responses were seen only in direct *ex vivo* or after irrelevant-peptide expansion, many responses such as SARS1, SARS2, EBV16, Flu1, and CMV27 were common across all conditions.

**Figure 2:**
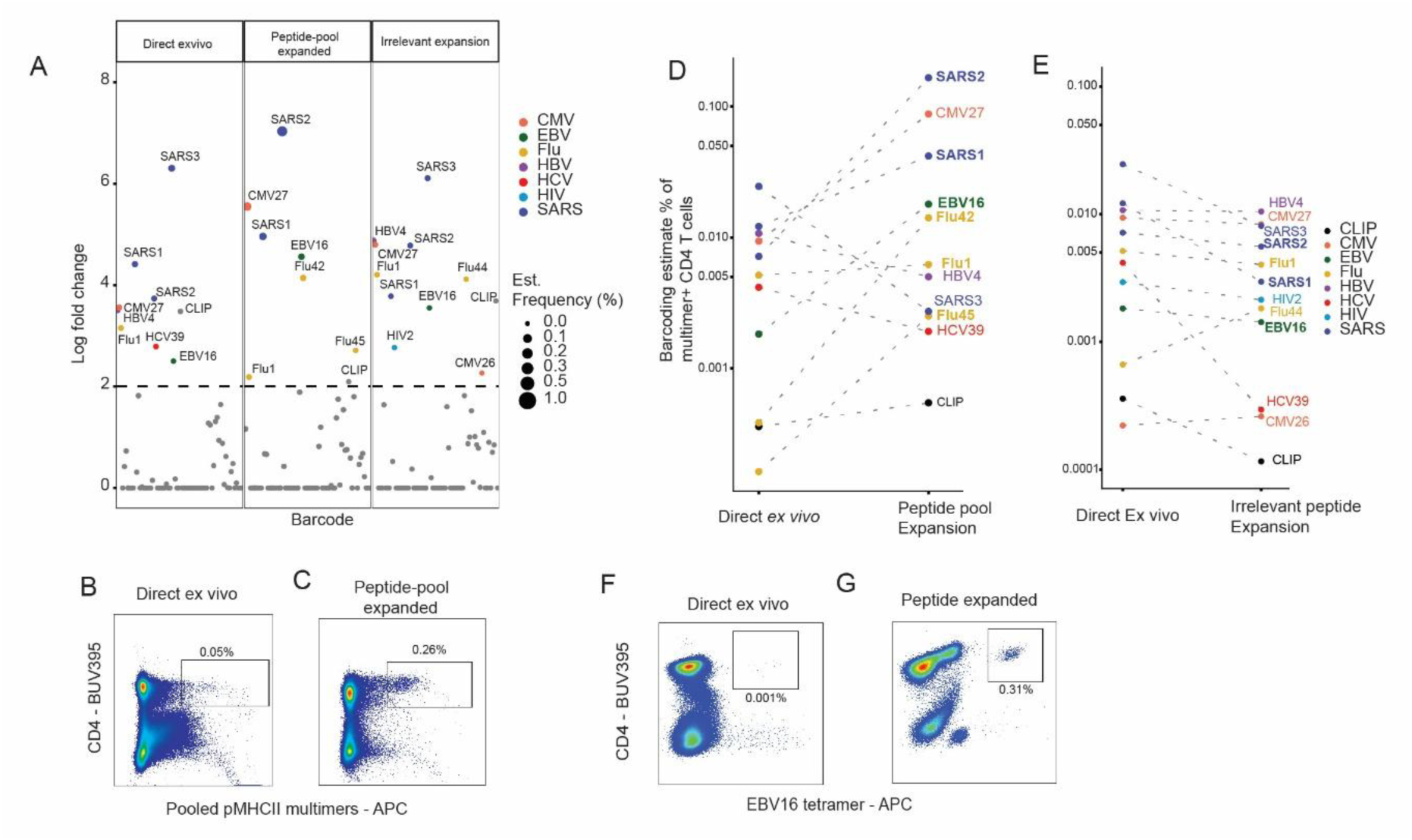
Detecting low-frequency virus-specific CD4 T-cell responses by peptide-pool expansion using barcoded-pMHCII multimers: A) Detection of responses in participant BC408, with a panel of 70 barcode-labelled pMHCII multimers (21 viral peptides + 49 irrelevant peptides). Three conditions are shown: Direct *Ex vivo* PBMC, PBMC expanded with the 21 viral-peptide pool (Peptide pool expansion) and PBMC expanded with an irrelevant HIV1 peptide (Irrelevant expansion). All significant pMHCII responses are colored and marked by the virus they are derived from. The dotted line represents significant enrichment threshold (logFC>2 and p<0.001). B-C) Flow plots showing CD4 T cells from *ex vivo* BC408 (B) and BC408 expanded with 21 viral-peptide pool (C) screened with 70 barcode-labelled pMHCII multimers (21 viral peptides + 49 irrelevant peptides). D) Change in percentage (%) of multimer-positive CD4 T cells comparing Direct *ex vivo* with Peptide-pool expanded BC408 sample. The peptide responses bolded were validated with tetramer stains on Direct ex vivo cells. The responses are colored as per the virus they are derived from. E) Shows the change in barcoding estimated percentage (%) of multimer positive CD4 T cells comparing Direct ex vivo with irrelevant-peptide expanded BC408 sample. The peptide responses bolded were validated with tetramer stains on Direct ex vivo cells. The responses are colored as per the virus they are derived from. F-G) Flow plots showing CD4 T cells from *ex vivo* BC408 (F) and BC408 expanded with EBV16 peptide (G) stained with EBV16 tetramers.

Following expansion with the peptide pool, the overall frequency of pMHCII multimer-binding cells increased notably from 0.08% to 0.3% of the CD4 T cell compartment (**Figure 2B-2C**). Correspondingly, most specific T cell responses showed increased estimated frequencies of multimer-positive CD4 T cells compared to the direct *ex vivo* sample, notably SARS2, CMV27, SARS1, EBV16, Flu42, and Flu45, suggesting that these pre-existing, antigen-specific responses were expanded in the presence of antigenic peptide (**Figure 2D**). In contrast, no T cell expansion was observed with irrelevant peptide, compared to the ex-vivo condition (**Figure 2E**).

To validate the responses detected within the HD1 screening, we performed direct tetramer staining with 7 pMHCII complexes, that were all seen as significant in peptide-pool expanded samples, of which 6 of them were validated by dextran multimer stains in direct *ex vivo* sample and individual peptide stimulation and expansion (**Supplementary figure 2**). We additionally also selected 4 peptides-MHCII for which T cell recognition was not confirmed in the peptide-stimulated sample (HBV4, SARS3, Flu44 and HIV2), and T cell recognition to these was not observed by individual tetramer staining (Supplementary figure 2). EBV16-specific CD4 T cells were seen at a frequency of 0.001% in direct *ex vivo* sample, which also showed peptide-specific barcode enrichment in *ex vivo* sample, confirming that the barcode-labelled pMHCII methodology possesses high sensitivity and can capture low-frequency virus-specific CD4 T cells at frequencies as low as 0.001% (**Figure 2F and 2G**). This suggests the detection of antigen-specific CD4 T cells depends also on the affinity of the peptide to the respective CD4 T cells, as some can be detected with better clarity than the others. Yet, the barcoded-MHCII multimers when applied on direct *ex vivo* and peptide-pool expanded samples showed that a combination of the two assays provides higher certainty of the actual antigen-specific CD4 T cells being present.

### DNA-barcoded pMHCII multimers can detect viral CD4 T cells in an expanded viral peptide panel on healthy donors with HLAs DRB1*01:01 and DRB1*04:01

To assess the broader capability of barcoded-pMHCII multimers in profiling CD4 T-cell responses, we expanded our library to a larger panel of 96 virus-derived peptides (**Figure 3A**) presented by HLAs DRB1*01:01 and DRB1*04:01. This comprehensive panel included epitopes from Influenza (Flu, n=41), Epstein-Barr Virus (EBV, n=25), Human Cytomegalovirus (CMV, n=23), Hepatitis B Virus (HBV, n=4), SARS-CoV-2 (n=2) and SARS-CoV-1 (n=1), and HCV39 (**Supplementary table 2**). The peptides were a combination of already characterized epitopes from IEDB and NetMHCIIPan predicted HLA-DRB1*01:01 and DRB1*04:01 peptide ligands for Flu, EBV and CMV against antigens hemagglutinin, nuclear antigen, and 65kDa phosphoprotein respectively. DNA-barcoded pMHCII multimer panel was generated for the viral peptides, which was then used to screen for virus-specific CD4 T-cell responses across a cohort of 8 healthy donors (6 donors for DRB1*01:01 and 2 donors for DRB1*04:01) as both direct *ex vivo* and peptide-pool expanded samples.

**Figure 3:**
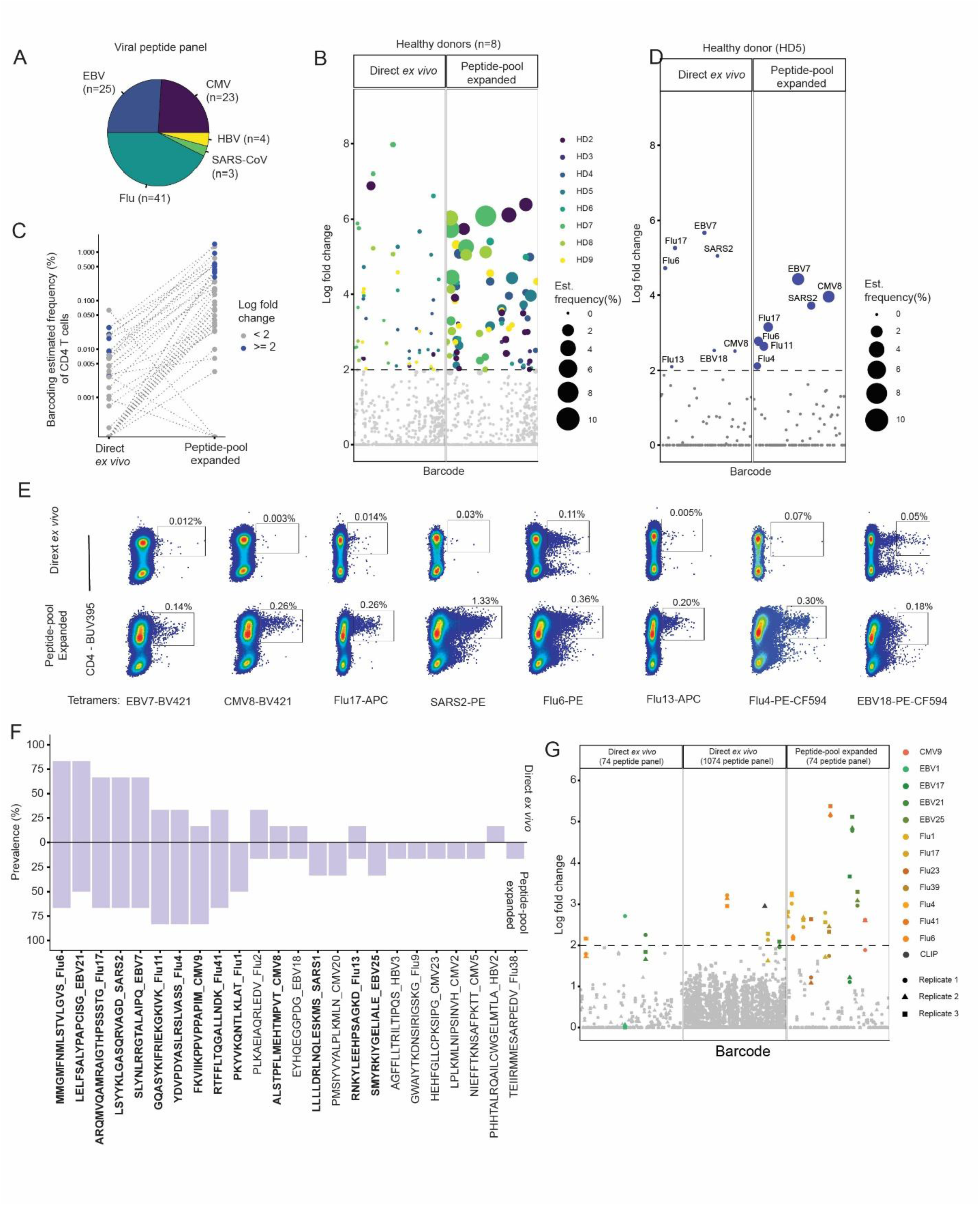
Application of DNA-barcoded MHCII multimer for detecting common virus-specific CD4 T cell responses in healthy individuals: A) Pie chart showing the distribution of a new panel of viral peptides (n=96) used for screening for virus-specific CD4 T cells in healthy donors. B) Viral-pMHCII responses within the sorted multimer+ CD4 T cell population, in 8 healthy donor samples, using the barcode-labelled-pMHCII multimers with the 96 virus derived peptides. The samples were screened as both direct *ex vivo* (triangles) and peptide-pool expanded (circles) samples. Each healthy donor is marked with individual colors. C) Change in barcoding estimated percentage (%) of multimer-positive CD4 T cells comparing Direct *ex vivo* with peptide-pool expanded for all participants (n=8). Dotted lines mark the respective response in Direct *ex vivo* and peptide-expanded sample in the same donor. The responses are colored blue and grey if their log fold change (logFC) value was >=2 or <2 respectively. D) Shows viral-pMHCII responses within the sorted Multimer+ CD4 T cell population for PBMC HD4, both in direct *ex vivo* (triangle) and peptide-pool expanded (circles). The size represents the estimated frequency of the peptide-specific CD4 T cells. E) Flow plots showing tetramer stains validating the responses detected and shown for HD4 for both direct *ex vivo* and peptide-pool expanded cells. F) Prevalence plot showing the prevalence of DRB1*01:01 restricted peptides in 6 samples plotted as % on y axis, and the peptides on x axis. The top segment represents prevalence in direct *ex vivo* and bottom segment represents prevalence in peptide-pool expanded samples. Peptide responses which could be validated by conventional tetramer stains in at least one PBMC are bolded. G) Plot showing the reproducibility of viral response detection in PBMC HD1. Each barcode enrichment is shown as log-fold-change and the dotted line represents the threshold for significant responses (logFC >2 and p<0.001). Responses identified as significant in at least one of replicates is colored as per the peptides and the replicates are shown in circles, triangle, and square for replicate 1, 2, and 3 respectively. The plots from left to right are direct *ex vivo* sample screened with 74 barcoded-pMHCII multimers, direct *ex vivo* sample screened with 1082 barcoded-pMHCII multimers (1000 DRB1*01:01-CLIP as irrelevant peptides), and the peptide-pool expanded sample screened with 82 barcoded-pMHCII multimers where all pMHCII have unique barcodes.

We observed multiple viral specific barcode enrichments in both direct *ex vivo* and peptide-pool expanded samples across all donors (**Figure 3B**). Importantly, these signal were enhanced following peptide pooled expansion, in 70 of 73 cases, indicating that the T cell recognition observed directly *ex-vivo* was indeed associated with true T cell responses (**Figure 3C**).

To further validate the presence to the detected T cell populations, using donor HD5 as example (**Figure 3D**) T cell recognitions observed using the barcoded pMHCII reagents, was confirmed with individual tetramer staining on both direct *ex vivo* and peptide–pool expanded cells. All responses observed was validated with tetramer staining (**Figure 3E**), although except the EBV18-MHCII complex, which provides significant background staining, possibly resulting in the loss of signal in the barcode-pMHCII screen in the peptide-expanded sample.

Next, we assessed the prevalence of T cell recognition across all tested DRB1*01:01 restricted healthy donors, in both direct ex-vivo evaluated samples and peptide-expanded samples, respectively (**Figure 3F**). The peptide-MHCII binding that was validated by pMHCII-tetramers in at least one of the heathy donors were bolded (**Figure 3F, Supplementary figure 3**). This reveals a number of epitopes showing high prevalence in both direct *ex vivo* and peptide-pool expanded samples, including Flu6, EBV21, Flu17, SARS2, and EBV7. Few others showed lower prevalence in *ex vivo*, whereas they had higher prevalence in peptide-pool expanded samples, such as Flu11, Flu14, CMV9 and Flu41, indicating that these are generally of low frequency challenging the detection in direct *ex-vivo*. Some epitopes such as Flu1, EBV25, and SARS1 could not be detected in direct *ex vivo* samples, but were detected in peptide-pool expanded samples. The prevalence for DRB1*04:01 restricted peptides were not plotted as there were only 2 healthy donors.

Finally, we wanted to address feasibility of using large libraries of pMHCII to screen for T cell recognition. Therefore, we extended the 74 DRB1*01:01 viral pMHCII-multimer panel with 1000 irrelevant pMHCII (DRB1*01:01-CLIP) each with unique DNA barcodes and compared the results when screening HD2 PBMC (**Figure 3G**). The 74 viral pMHCII-multimer panel showed peptide-specific barcode enrichment towards EBV21 and Flu6, which were also detected when the panel was increased with 1000 irrelevant pMHCII suggesting that the peptide-specific CD4 T cells that can be detected will be detected even with increasing the barcoded-pMHCII multimer panel size. Further, to test the reproducibility of the DNA-barcoded pMHCII multimers across direct *ex vivo* and peptide-pool expanded samples, we performed three independent pMHCII multimer screenings and observed good consistency comparing the data (**Figure 3G**). In addition, most of the barcode-enriched viral responses could be validated both direct *ex vivo* and after peptide –pool-expansion (**Supplementary figure 4**).

Interestingly, validation of the healthy donors using conventional tetramer revealed cross-reactive responses in three of the PBMC (**Supplementary figure 5**). We observed a clear double-positive CD4 T cell population that co-stained with Flu41 (RTFFLTQGALLNDK) and SARS1 (SARS-CoV 2 - LLLLDRLNQLESKMS) in two of the healthy donors. In one of them (HD2, we could confirm the cross-reactivity as both direct ex vivo and peptide-pool expanded cells. The Flu41 peptide is a novel epitope which is not documented in IEDB, but part of a previously-reported 17-mer peptide^29^. We also observed cross reactivity between EBV21 (LELFSALYPAPCISG) and CMV9 (FKVIIKPPVPPAPIM) in one healthy donor, but could not validate it by peptide expansion due to lack of PBMC material.

### Application of DNA-barcoded pMHCII multimers to screen acute HCV patients revealed novel HCV epitopes restricted to DRB1*01:01 and DRB1*04:01

We applied the DNA-barcoded pMHCII multimer screening approach on a cohort of acute HCV patients restricted to DRB1*01:01 (n=6) and DRB1*04:01 (n=8). To achieve this, we first assessed cohort of participants with acute HCV infection, restricted to DRB1*01:01 and DRB1*04:01, who were pre-screened and known to have at least one viral-specific CD4 T cell response detected by tetramer or Intracellular Cytokine Staining (ICS) assays (14 participants with a known HCV-specific response, of which 3 also had a known EBV- or CMV-specific response, and 2 participants with only a known EBV- or CMV-specific response). Forty-five 20mer HCV peptides that were previously found to be positive in ICS in participants with DRB1*01:01 and/or DRB1*04:01 alleles were assessed in NetMHCIIpan to determine the DRB1*01:01 and/or DRB1*04:01 binding sequences. Based on the elution rank and the unique binding cores we generated 53 peptides (Supplementary table 3) covering all structural and non-structural proteins from the HCV polyprotein except for p7 (**Figure 4A**).

**Figure 4:**
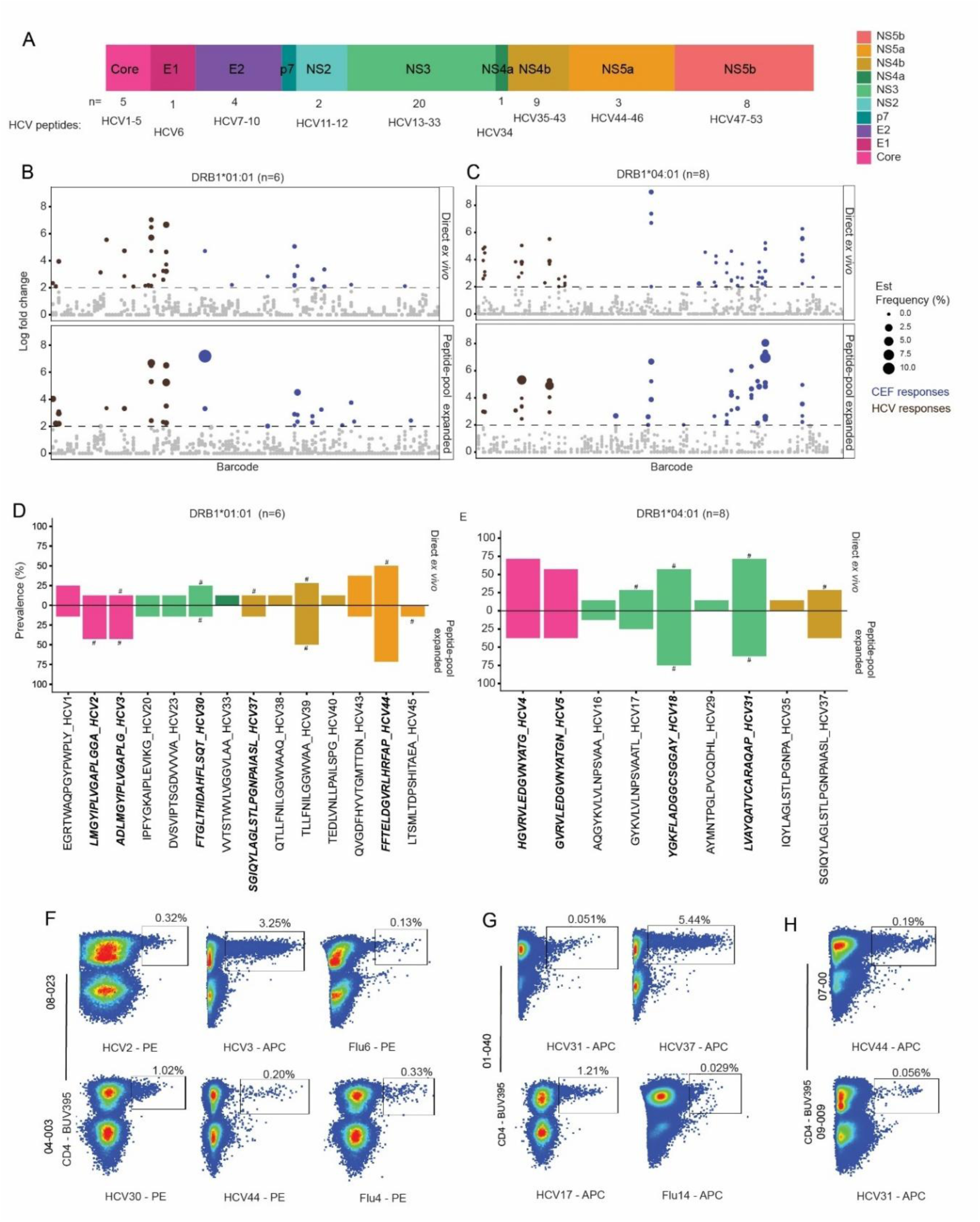
Using DNA-barcoded pMHCII multimers to screen and detect HCV-specific CD4 T cells in participants with HCV infection. A) The HCV viral genome and the respective peptides derived from the color-coded proteins. The size of the genome segment represents the number of predicted peptides from that segment. B-C) pMHCII responses within the sorted Multimer+ CD4 T-cell population. DRB1*01:01(B) participants (n=6) where screened with a panel of 122 barcode-labelled pMHCII multimers and DRB1*04:01 (C) participants (n=8) were screened with a panel of 108 barcode-labelled pMHCII multimers. The participants were screened both directly *ex vivo* (top) and after peptide-pool expansion (bottom). Common viral responses are shown in blue and HCV responses are shown in brown. The size of the dots represents the estimated population frequency. D-E) Displays the prevalence of the HCV-derived peptides from the multimer screens. Direct ex vivo is shown on the top and peptide expanded shown below. DRB1*01:01 is shown in (D) and DRB1*04:01 is shown in (E). The color of the bar represents the segment of the virus the peptide derives from (A). Bars with a # were validated with tetramer stains within the participants and text in bold have been validated with tetramer stains in external participants. F) Example tetramer staining flow plots validating HCV- and CEF-specific responses in DRB1*01:01 participants after peptide-specific expansion. G) Tetramer staining flow plots validating HCV- and CEF-specific responses in DRB1*04:01 participants after peptide-specific expansion. H) Tetramer staining flow plots showing HCV31- and HCV44-specific CD4 T cell detection in participants 09-009 and 07-00 (Patients not within the HCV cohort).

Combining this DNA-barcoded HCV-specific library with the virus library (**Figure 3A**), we screened 6 and 8 participants with DRB1*01:01 and DRB1*04:01 respectively, using both direct *ex vivo* and peptide-pool expanded samples. T cell-recognition to multiple epitopes were detected, 22 HCV-derived pMHCII and 29 CEF-derived pMHCII. From the HCV pool, 9 of these are potentially novel MHCII-presented epitopes. We observed enhanced signal of HCV-specific population following peptide-pool-expansion, indicating functional T cell reactivity (**Figure 4B-C**).

For all HCV-specific peptides with detectable T cell recognition we calculated the prevalence (%) in direct *ex vivo* and peptide-pool expanded samples for across both DRB1*01:01 restricted (**Figure 4D**) and DRB1*04:01 restricted (**Figure 4E**) cohorts. Several T cell populations were additionally validated by conventional tetramer staining using peptide-specific expansion samples from the respective patient material (**Figure 4F, Supplementary figure 6, 7, and 8**, and marked by hash mark in Figure 4D and 4E). In the HCV cohort, we also validated some of the virus-specific responses using conventional tetramers (**Figure 4F and 4G**). In total, we detected T cell recognition towards 12 HCV epitopes, including 9 epitopes not previously used in multimer assays of which HCV2 and HCV3; HCV4 and HCV5 had similar binding core and have only two amino acid difference in the peptide sequence. HCV2, HCV30, and HCV31 were previously seen positive for ICS studies, but not restricted to any HLA. HCV3, HCV5, and HCV18 were not documented prior to this work. The HCV-specific epitopes that were detected in this pMHCII multimer screens were also assessed in 11 additional participants where we saw peptide-specific CD4 T cells in the tetramer stains, confirming the broader use of these epitopes to screen HCV patients(**Figure 4H**).

Across the different peptides that were identified for DRB1*01:01 restriction, 2 were derived from the core protein forming the viral nucleocapsid and the rest were from the non-structural region of the virus genome, mostly NS3, NS4b, and NS5a. For DRB1*04:01 restricted peptides, we could identify two peptides in the core protein, but could not be validated on the cohort although it was validated in external participants (**Supplementary figure 9**); and the rest were in the non-structural portion of the HCV virus.

## Discussion

Detection and characterization of antigen-specific CD4⁺ T cells has long been lagging behind equivalent efforts for CD8⁺ T cells ^30^. This gap stems from a combination of biological and technical challenges inherent to the MHCII system. Biophysically, the CD4-MHCII interaction is weaker than the CD8–MHCI interaction, and the TCR–pMHCII interaction is itself ∼4–5-fold weaker than TCR–pMHCI binding, as measured by surface plasmon resonance ^31,^35,^33^. These low-affinity interactions are compounded by the instability of recombinant MHCII heterodimers, limiting the production of reliable soluble reagents. Click or tap here to enter text.Beyond reagent limitations, antigen-specific CD4⁺ T cells circulate at very low frequencies in peripheral blood, often below the detection threshold of direct ex vivo assays ^34^. To overcome this, many studies have relied on in vitro expansion which significantly alter T cell phenotype and function and selectively enrich only cells capable of proliferation or defined marker upregulation ^35^. Alternatively, functional assays such as ELISpot, intracellular cytokine staining, and cytokine release assays require large amounts of cellular material and either of them use pooled peptides that sacrifice individual epitope-level resolution ^36^.^40^ As a result, it has remained particularly difficult to discover novel CD4⁺ T cell epitopes the resolution of individual antigenic determinants across large antigen libraries.

Here, we present a DNA-barcoded pMHCII multimer technology that directly addresses the limitation by enabling simultaneous screening using large libraries of up to 1000 peptide–MHCII complexes with unique DNA barcodes — within a single sample. Unlike conventional functional assays or pooled peptide approaches, this platform identifies the peptide–MHCII specificity of endogenous CD4⁺ T cells at epitope resolution. The scalability of this approach represents a fundamental advance over pre-existing methods, which are constrained to testing one peptide specificity per well or reporting only aggregate reactivity to peptide pools.

To address the weak TCR–MHCII interaction, we employed dextran-based pMHCII multimers, which offer increased valency. Whether dextran-based multimers outperform conventional tetramers in this regard, however, remains to be directly evaluated^37^. Dextran scaffolds have previously demonstrated utility in detecting low-affinity MHCI–TCR interactions, and their increased valency equivalently strengthens MHCII–TCR interactions, enabling detection of antigen-specific CD4⁺ T cells at frequencies as low as 0.001% direct *ex vivo* in some donors and certain specificity. To further enhance the CD4–MHCII interaction itself, affinity-enhanced MHCII reagents incorporating two point mutations at the CD4-contacting β2 domain of HLA-DRB1*01:01 have been developed, improving CD4 affinity approximately 200–1000-fold (KD∼∼5–9 µM), bringing it into the same range as the CD8–MHCI interaction ^38^. Integration of such affinity-enhanced reagents with the DNA-barcoded multimer platform holds considerable promise for extending detection to CD4⁺ T cells with even lower TCR affinity.

The challenge of low circulating CD4⁺ T cell frequencies was addressed by integrating peptide-stimulated in vitro expansion prior to barcoded pMHCII multimer screening. This strategy selectively enriches antigen-specific CD4⁺ T cells, allowing reliable detection of T cell populations that fall below the direct ex vivo detection threshold. We observed a good overlap in peptide-specific barcode enrichment (38 epitopes across 8 healthy individuals)tection of antigen specific T cell populations between direct ex vivo and peptide-stimulated conditions, confirming that expansion does support in detecting the antigen-specific CD4 T cells with higher certainity. Peptides uniquely detected after stimulation are likely low frequent T cell populations undetectable directly ex-vivo. This two-stage approach therefore preserves the epitope-level resolution of the multimer platform while extending its sensitivity. Although, asymmetric clonal expansion should be considered when using peptide-pooled stimulation and expansion, especially when number of peptides are scaled up ^39^. To ensure that fast-expanding clonotypes do not outcompete the smaller population, we kept the in-vitro expansion only until Day-9.

Click or tap here to enter text.Click or tap here to enter text.Click or tap here to enter text.Click or tap here to enter text.Click or tap here to enter text.Click or tap here to enter text.Non-specific MHCII binding was tested through the inclusion of pMHCII-CLIP with each unique DNA barcodes as internal control in every screening experiment. CLIP barcodes were enriched in 3 out of 24 donors screened, confirming that background signal cannot be entirely eliminated but can be systematically identified and excluded. Critically, when the number of DRB1*01:01-CLIP barcodes was scaled up to 1000, only one corresponding barcode was enriched, demonstrating that background-staining with MHCII reagents has very little impact in detection of antigen-specific CD4 T cells when using DNA-barcoded pMHCII multimers.

The platform’s capacity to detect CD4⁺ T cell responses was validated across multiple viral antigens in both healthy and diseased individuals. In the context of HCV infection, where 70–80% of patients fail to clear the virus and develop chronic infection, we identified 8 novel CD4⁺ T cell epitopes. Several of these epitopes were validated in patients outside the discovery cohort, demonstrating that they are recognized broadly across individuals and suggesting their potential utility as vaccine candidates ^19^. We also detected HIV-specific barcode enrichment in donors stimulated with an irrelevant peptide, which may reflect cross-reactive CD4⁺ T cells primed by microbiome-derived peptides sharing 8–12 amino acids with HIV sequences, an observation consistent with prior reports of HIV-specific responses in unexposed individuals ^40,41^.

The development of this DNA-barcoded pMHCII multimer technology represents a significant methodological advance in T cell immunology, providing a capacity to identify CD4⁺ T cell responses at the level of individual epitopes and across large antigen libraries simultaneously. Of particular significance is the potential to simultaneously interrogate CD4⁺ and CD8⁺ T cell specificities within the same donor, enabling direct characterization of antigen and epitope overlap between the two compartments — a question that has remained largely unaddressed outside of defined model antigens. Such resolution is of considerable importance for rational vaccine development, as effective subunit and epitope-based vaccines require a thorough understanding of which antigenic determinants are immunologically relevant and how broadly they are recognized across individuals.

As this technology evolves — with advances in affinity-enhanced reagents, broader MHC allele coverage, and integration with single-cell genomic platforms — it holds considerable promise as an important tool for understanding human T cell immunity and translating these insights into more effective and precisely targeted immunological therapies. This novel tool shows strong potential, and future integration of larger prediction datasets and additional HLA alleles may further enhance epitope discovery. Additionally, future incorporation of single-cell sequencing, TCR repertoire analysis, and multimodal immune profiling could further expand the utility of this approach. In summary, we establish DNA-barcoded pMHCII multimers as a robust and scalable platform for large-scale identification of antigen-specific CD4 T cells across extensive peptide libraries, facilitates discovery of previously unrecognized epitopes, and provides a framework for comprehensive immune monitoring in infectious diseases, cancer, autoimmunity, and vaccine development and its potential utility for future vaccine design and immunotherapeutic strategies.

## Acknowledgement

We would like to thank United States, Department of Health and Human Services, National Institutes of Health, National Institute of Mental Health for sharing the HEK-293 cell line expressing human DRB1*01:01 and DRB1*04:01. We would like to thank Mauricio Calvo-Calle and Aniuska R. Becerra Artiles for sharing the protocol for CD4 T-cell expansion.

## Author contribution

S.R.H. conceived the idea. Y.B., T.T., S.D., G.M.L., and S.R.H. designed the study. Y.B., T.T., L.L., L.S., and W.W.K. contributed to reagent production. Y.B., T.T., S.K.S., and S.D. performed experiments. Y.B., T.T., S.D., G.M.L, S.R.H performed data analysis and interpretation. Y.B., S.K.S., and S.R.H. prepared figures. Y.B., S.K.S., and S.D. contributed to sample collection. Y.B., T.T., S.K.S., S.D., and S.R.H. wrote the manuscript. S.R.H supervised the study and verified the underlying data of this manuscript.

## Methods

### Plasmid isolation

The plasmid to be isolated was transformed to one shot Stbl3 chemical competent cells (C7373-03 Thermofisher) per the manufacturer’s instructions at 42°C for 45 seconds, plated on an LB agar plate containing carbenicillin (100µg/mL) and incubated at 37°C incubator. The next day, a single colony was inoculated into a conical flask containing 200mL of LB media with carbenicillin (100µg/mL) for plasmid isolation. Plasmid was isolated the next day using a Qiagen Plasmid plus Midi kit (Qiagen, 12945). The plasmid isolated was then measured using Nanodrop ND-1000.

### TCR expressing supT1 cell line generation

TCR sequence which was previously documented ^42^ and showed binding to HA_306-318_ (Flu1) loaded DRB1*01:01 and DRB1*04:01 – HA1.7 was obtained from PDB ID-1J8H^43 44^ and synthetically constructed with codon-optimization to be expressed in HEK-293 cells. HEK293T (CRL-3216) was thawed and cultured at least twice before transfection in RPMI-1640 + 10% FBS by carefully using Trypsin-EDTA to release the adhered cells in a T-75 culture flask. The cells were seeded to ensure approximately 50% confluency in a T-75 flask 24 hours before transfection. The cells were then transfected with 15µg of the vector-of-interest and the packaging plasmid (REV, Gag/pol, and VSV glycoprotein) using Lipofectamine 3000 (Thermofisher L3000008) per manufacturer’s instructions. The virus was collected 24 and 48 hours after transfection and concentrated using a Lenti-X concentrator (Clontech 631232) as per the manufacturer’s instructions. The concentrated virus was then resuspended in 200µL PBS and frozen in aliquots of 50µL at −80°C. SupT1 cells (CRL-1942) was cultured in RPMI-1640 +10% FBS + 1% Penicillin-Streptomycin (Thermofisher 15140122) and passaged at least twice before transduction. 1 × 10^6^ cells were seeded into a single well of a 24-well plate. To this, one aliquot (50µL) of the frozen virus is added by diluting it with 200µL RPMI-1640. The cells were mixed well and incubated in a 37°C, 5% CO2 incubator at 85% humidity for 48 hours. The cells were then scaled up to 3mL in a 6-well plate. At this stage, the cells were analyzed by flow cytometry for expression of GFP (Supplementary figure 1). The transduced cells were scaled up and frozen in RPMI-1640+10%FBS+5% DMSO in liquid nitrogen.

### MHCII-Construct, expression, and purification

Soluble MHC class II DNA constructs were generated by taking the alpha- and beta-subunit sequences from NIH ^45^, and codon-optimized to be expressed in HEK293 cells, and the two ectodomains were separated by a P2A ribosomal skipping sequence and under a control of CMV promoter (pCDNA 3.4 plasmid, Thermofisher). MHCII was expressed using the Expi293 expression system (Thermofisher A14635) per the manufacturer’s instructions. Expi293F cells were cultured in Expi293 expression medium (Thermofisher A1435102) by maintaining an initial cell density of 0.3-0.5 × 10^6^ cells/mL in 125mL sterile shake flask (Corning 431143) and incubated in 37°C, 8% CO_2,_ 85% humidity and 125 rpm. The cells were passaged at least thrice before taking it forward for transient expression study. On Day 0, plasmid DNA (1µg/mL) was transfected to Expi293 culture using the Expifectamine transfection reagent as per manufacturer’s instructions. Transfection kit enhancers 1 and 2 were added on Day 1. Cultures were harvested on Day 6 of transfection by centrifuging at 16000 x g for 45 min and sodium azide was added to get a final concentration of 0.02%.

When the soluble MHCII was expressed using NIH cell line, the cell line was cultured in complete media (RPMI-1640 + 10% FBS) for a period of 7 days and then the media was changed to Freestyle 293 serum-free media (ThermoFisher, 12338018). The cells were cultured in serum-free media for 10-14 days, after which the supernatant was harvested.

The harvested supernatant in both cases was then filtered through a 0.22µm regenerated cellulose membrane. The filtered supernatant was then supplemented with 40mM Tris-HCl pH 8.0, 300mM NaCl, 23% glycerol, and 1mM PMSF prior to protein purification. The harvested supernatant was concentrated using Amicon Ultracel 30K MWCO regenerated cellulose devices and buffer-exchanged with PBS pH 7. The buffer-exchanged protein supernatant was biotinylated with BirA biotin-protein ligase bulk reaction kit (5.5 µg BirA per 50 nmol of MHCII protein) overnight at room temperature. Digested and biotinylated MHCII protein was finally purified via size exclusion chromatography using a Superdex 200 10/300 GL increase column (Cytiva) equilibrated with 1X PBS. Fractions corresponding to the soluble MHCII (typically centered at 13 mL retention volume) were identified via SDS-PAGE, pooled, and concentrated using Amicon Ultracel 30K MWCO regenerated cellulose devices to ∼2-3 mg/mL. Aliquots were stored at −80°C.

### HLA-DM: Expression and Purification

A soluble HLA-DM (HLA-sDM) expressing cell line was the courtesy of Elizabeth Mellins (Stanford University) ^46^. The construct had been generated in Drosophila S2 cells in which the transmembrane and cytoplasmic domains of DM-α and DM-β were truncated and replaced with FLAG and simian virus 40 (SV40) large T-antigen epitope tags, respectively as per Sloan et. al.^47^ The soluble HLA-DM S2 cell line was cultured in Sf900 II SFM (Thermofisher 21012026) in the presence of 10% FBS and 1µg/mL Geneticin (Thermofisher 10131035) at 28°C until the cell count was >5 × 10^6^ cells/mL. At this point, it was changed to gentle shaking at 120rpm and further, the FBS was weaned out from 10% to 0% slowly with 2 passages at each concentration of FBS. When the cells were in serum-free condition, the cells were then seeded at 6-8 × 10^6^ cells/mL and induced with 1mM CuSO4 solution for 7 days, after which the culture supernatant was harvested by centrifuging at 3500 x g for 45 min. The supernatant was removed, and sodium azide was added to get a final concentration of 0.02%. The supernatant was then filtered through a 0.22µm regenerated cellulose membrane. HLA-sDM was then purified by affinity chromatography using AntiFLAG M2 affinity gel (Sigma-Aldrich A2220) followed by purification using Superdex 200 10/300 GL increase column (Cytiva) equilibrated with 1X PBS. Fractions corresponding to the soluble HLA-DM were identified via SDS-PAGE, pooled, and concentrated using Amicon Ultracel 30K MWCO regenerated cellulose devices to ∼1 mg/mL. Aliquots were stored at −80°C.

### Peptide prediction

All viral peptides (Human CMV, EBV, Flu, HBV, SARS-CoV-1, and SARS-CoV-2) other than HIV were obtained either from IEDB or predicted using NetMHCIIpan ^48^. The predicted peptides were obtained from antigens of common viruses – Hemaglutinin for Flu, nuclear antigen for EBV and and 65kDa phosphoprotein for human CMV. The peptides were selected based on unique binding core and their Rank score (<10). In total we had 23 peptides for human CMV, 25 peptides for EBV, 40 peptides for Flu, 4 peptides for HBV, 1 peptide for SARS-CoV-1, and 2 peptides for SARS-CoV-2 (Supplementary table 2). The two HIV derived peptides –HIV1 – FRKQNPDIVIYQYMDDLYVG restricted to DRB1*01:01 with epitope origin of HIV reverse transcriptase 171-190^49 50^ and HIV2 – ACQGVGGPGHKARVLA as an irrelevant peptide^51^ were not sourced from IEDB. HCV peptides were predicted from 20mer peptides which were used to study the CD4 T cell reactivity in HCV patient samples ^52^. The 20mer peptides were then trimmed using NetMHCIIpan, and chosen based on Rank score (<5) and unique binding core.

### Peptide exchange

Soluble MHCII protein was incubated with 3C protease in cleavage buffer (Tris-HCl pH 7.5 and NaCl,) overnight at 4°C. The next day, protease-cleaved MHCII protein (10 µM) was incubated with 100 µM of peptide of interest in an exchange buffer (100 mM sodium acetate pH 5.2, 50 mM NaCl, and 5 µM HLA-sDM) for 18 h at 37°C ^53^. After incubation, the reaction mixture was centrifuged at 3300 x g for 10 min to remove aggregates, and the supernatant was transferred and neutralized with 2X PBS final concentration. Exchanged peptide-MHCII (pMHCII) monomers were then freshly conjugated with fluorochrome-labelled streptavidin or stored at −80°C until further use.

### DNA-barcoded multimer library preparation

The DNA-barcoded multimer library was prepared using the method developed by Bentzen et al ^54^. Different oligos were combined to generate unique barcodes, with each barcode containing a 5’ biotinylated unique DNA sequence. These barcodes were then attached to streptavidin-conjugated dextran (Fina BioSolutions, Rockville, MD, USA) coupled to allophycocyanin (APC) or phycoerythrin (PE) fluorophore by incubating them at 4°C for 30 min to generate a DNA-barcode-dextran library containing 1074 unique barcodes. DNA-barcoded multimer libraries were then generated by incubating the peptide-exchanged MHCII monomers to their corresponding DNA barcode-labelled dextrans at 4°C for 30 min, thus providing a DNA barcode-labelled peptide-MHCII (pMHCII) multimer.

### PBMC staining with DNA-barcoded pMHCII multimers

All participant PBMCs were thawed in prewarmed RPMI-1640 + 10% FCS + DNase (final concentration of 100µg/mL) and centrifuged at 400 x g for 5 min. If the PBMCs needed to be enriched for CD4 T cells, they were resuspended in RPMI-1640 + 10% FCS + DNase and incubated at room temperature for at least 15 minutes. Following this, the cells were normalized to 50 × 10^6 cells/mL and negatively enriched for CD4+ cells using EasySep Human T cell isolation kit (Cat. 19052) from Stemcell technologies. The enriched cells or directly thawed PBMC were then counted using Chemometec Nuclecounter and washed twice with barcode cytometry buffer (PBS with 0.5% BSA, 100 ug/ml herring DNA, 2 mM EDTA) before proceeding to staining. Cells were stained with 2µL of the MHCII-multimer at 37°C for 60 min and then stained with an antibody panel consisting of CD4-BUV395 (BD biosciences 563550), CD8-BV480 (BD biosciences 566121), and a dead cell marker (LIVE/DEAD fixable Near-IR; Invitrogen L10119). The plate was incubated on ice for 30 min. Cells were washed twice with barcode cytometry buffer (PBS with 0.5% BSA, 100 ug/ml herring DNA, 2 mM EDTA), fixed in 1% paraformaldehyde and acquired on a FACSAria flow cytometer instrument (AriaFusion, Becton Dickinson) as per the gating strategy in Supplementary figure 10. All CD4+, APC+ (pMHCII-multimer binding) cells were sorted (Supplementary figure 9) into pre-saturated eppendorf tubes with 2% Bovine serum albumin (BSA). The Eppendorf tubes are saturated with 2% BSA 1 hour before use, after which it is replaced with 100μl BCB. After sorting, the tubes are centrifuged for 10 min at 5000 × g. The supernatant was discarded with minimal residual volume, and the DNA barcodes, remaining in the cell pellet, were amplified using the Taq PCR Master Mix Kit (Qiagen, 201443) and sample-specific forward primer (serving as sample identifier) A-key. PCR-amplified DNA barcodes were purified using the QIAquick PCR Purification kit (Qiagen, 28104) and sequenced at PrimBio (USA) using the Ion Torrent PGM 314 or 316 chip (Life Technologies).

### DNA barcode amplification

Taq PCR Master Mix Kit (Qiagen, 201443) was used for amplification of DNA barcodes using 0.3 μM of appropriate forward and reverse primers comprising of adaptors (A-key and P1-key, respectively). PCR amplification was conducted on isolated cells (in <20 μL of buffer) or on the stored aliquot of the MHC multimer reagent pool (diluted 50,000× in the final PCR) used as baseline to determine the number of DNA barcode reads within a non-processed MHC multimer reagent library. Each sample was assigned a distinct sampleID embedded in the forward primer (A-Key). PCR was performed under the following conditions: 95 °C 10 min; 36 cycles: 95 °C 30 s, 60 °C 45 s, 72 °C 30 s; and 72 °C 4 min. The PCR products were purified with QIAquick PCR Purification kit (Qiagen, 28104) or QIAquick gel extraction kit (Qiagen, 28704). The amplified DNA barcodes were sequenced at using Nanopore sequencing.

### DNA-barcode sequence analysis and identification of pMHCII specificities

To process the sequencing data and automatically identify the barcode sequences, a specific software package was used, ‘Barracoda’ (https://services.healthtech.dtu.dk/service.php?Barracoda1.8). Fold changes in read counts mapped to a given sample relative to mean read counts mapped to triplicate baseline samples were estimated using normalization factors determined by the trimmed mean of M-values method. P values were calculated by comparing each experiment individually to the mean baseline sample reads using a negative binomial distribution with a fixed dispersion parameter set to 0.1 ^54^. The analyzed data was then wrangled in R and plotted to generate the respective figures. Barcoded-multimer pMHCII experiments were analyzed including a HLA-mismatch control. If any pMHCII-specific barcode enrichment in the HLA-mismatch control was observed, these pMHCII-specific barcodes were excluded from all the samples within that experiment. Significant barcode-enrichment of HBV1, HBV2, and HBV3 were removed from analysis for HD1 as they were enriched in HLA-mismatch control PBMC HD10 (Supplementary figure 11). For Figure 3B CLIP- and Flu36-specific barcode enrichment and for Figure 4B and 4C, HCV21-specific barcode enrichment was excluded due to high background staining. ^64, 65^

### Stimulation and expansion of peptide-specific CD4 T cells in PBMC

All PBMCs were thawed in prewarmed RPMI-1640 + 10% FCS + DNase (final concentration of 100µg/mL), and centrifuged at 400 x g for 5min. The cells were washed with RPMI-1640 + 10% FCS without DNase twice and then resuspended at a concentration of 2 × 10^6 cells/mL in expansion media - X-VIVO 15 media (Lonza, BE02-060Q) supplemented with 5% human serum (Sigma Aldrich Heat Inactivated H3667) and soluble IL-2 at 100U/mL (Preprotech). The resuspended cells were inoculated in a single well in a 24-well plate to which respective peptide/peptide pool was added to get a final concentration of 10µg/mL. The plates were then incubated at 37°C, 5% CO_2_. Expansion media with an IL-2 was added to the wells on Days 4 and 7 to get a final IL-2 concentration of 100U/mL. The cells were harvested on Day 9 and taken forward for staining.

### Tetramerization

Conventional tetramers were prepared by adding 18.1µL of streptavidin-fluorophore conjugate of 0.1mg/ml concentration per 100µL of peptide-exchanged MHCII monomer. It is then incubated on ice for 20 - 30 min, after which 25µM free biotin was added and incubated for another 20 min. Finally, 10x tetramer freezing buffer (50% glycerol+ 1% BSA) was added to get a final concentration of 1x and the tetramers are frozen in −20°C until further use.

### T cell staining

All PBMCs were thawed in prewarmed RPMI-1640 (Thermofisher, 11875093) + 10% FBS (Thermofisher, 10500064) + DNase (final concentration of 100µg/mL), and centrifuged at 400xg for 5min. Up to 5 million cells are pipetted into a well of a 96-well plate and the plate is centrifuged at 400 x g for 5 min and washed twice with FACS buffer (PBS pH 7.4 + 2% FBS). Cells were then stained with 1µL of the MHCII-streptavidin tetramer at 37°C for 60 min following which antibodies with live-dead marker was added. The plate was incubated on ice for 30 min. The cells were then washed with FACS buffer twice before being analyzed on flow cytometry. If the cells were not analyzed on flow cytometry, they were fixed with 1% PFA 1 hour to overnight, and washed twice with FACS buffer after fixing before analyzing on flow cytometry.

### Validation of HCV-specific responses

To validate HCV- and CEF-specific responses in participants from the HCV cohort, 5-20 million PBMCs per participant were used to generate short-term T-cell lines specific to each individual’s responses identified in the barcoded multimer screen. For each participant, peptides were grouped based on their barcode-estimated frequency to generate T-cell lines with a maximum of eight different peptides per culture flask. Peptide specific T-cell lines were generated as described previously ^55^. Briefly, PBMCs were thawed and rested for 2 h, after which they were stimulated with the peptides at 1 ug/mL for 1 h in R10 medium (RPMI 1640 (Sigma-Aldrich) supplemented with 10% FBS (Sigma-Aldrich), 10 mM HEPES buffer (Corning), 2 mM L-Glutamine (Corning), and 1% Penicillin-Streptomycin (Sigma-Aldrich)). Cells were then seeded at 1 M cells per mL of R10 and incubated at 37 degrees Celsius. After 1 day, 50 U/mL recombinant IL-2 (Sigma-Aldrich) was added to the media and fresh IL-2-containing media was added every other day. After 10-12 days, tetramer staining was performed as follows: cells were first stained with a viability dye (Life technologies), washed, and incubated with PE- and/or APC- and/or BV421-labelled MHC class II tetramers for 60 min at 37 degrees Celsius. The samples were gently resuspended every 20 min. After washing, populations were stained with surface antibodies for CD14, CD19, CD56, CD3, and CD4 for 30 min at 4 °C. Following a final wash, cells were resuspended in 2% paraformaldehyde (PFA) and acquired on a Bigfoot flow cytometer (Invitrogen). Data analysis was performed using FlowJo software, where the presence of singlet, lymphocyte, alive, CD14-, CD19-, CD56-, CD3+, CD4+, tetramer+ cells was assessed. After confirming responses in the participants that had been part of the barcoded multimer screen, PBMCs from an additional cohort of 11 participants (5x DRB01:01 and 6x DRB04:01) were used to generate T-cell lines for all novel HCV epitopes, with all HLA-matched reagents used for each participant.

## Supplementary Tables

**Supplementary Table 1:**
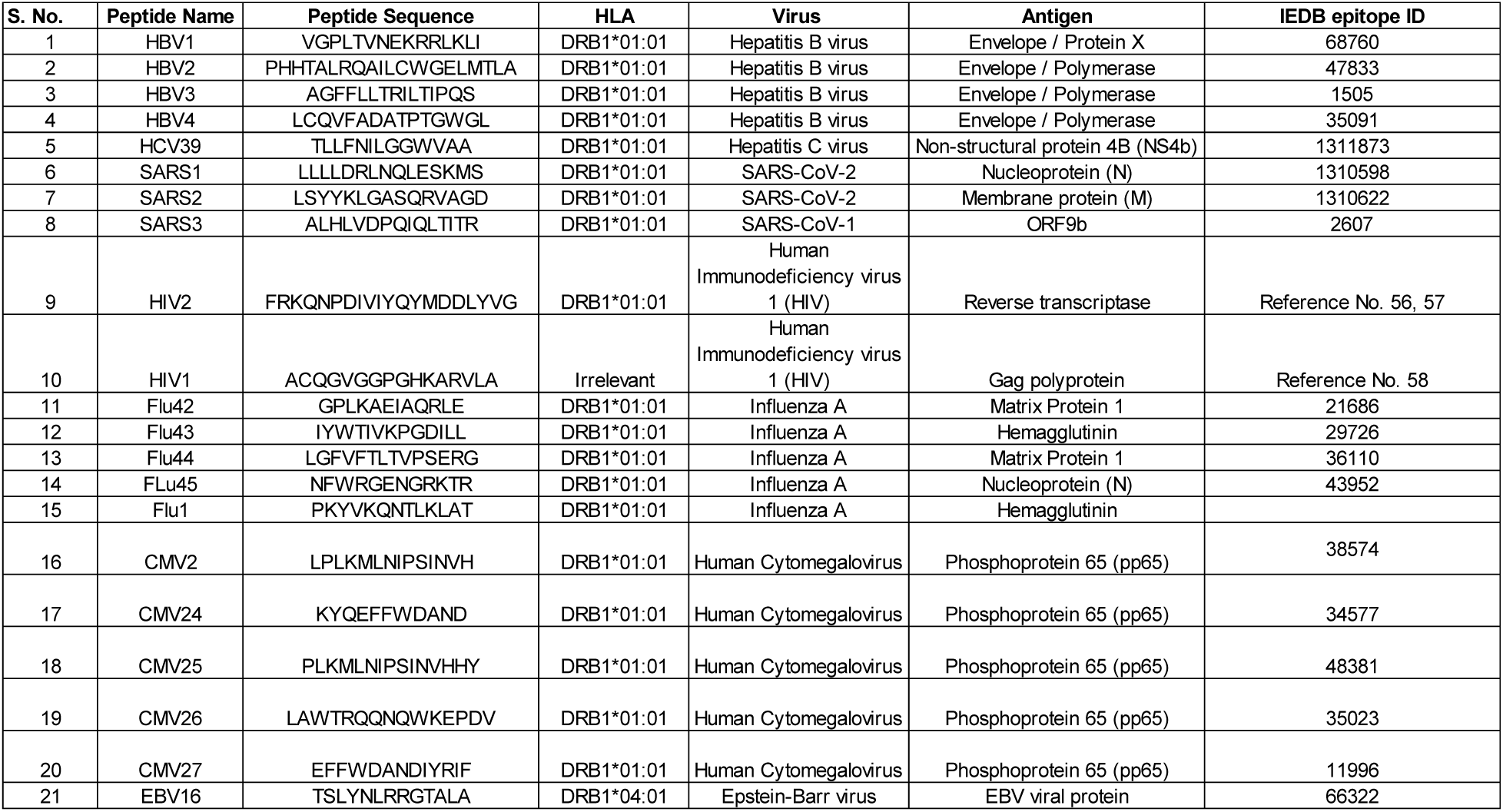
The table provides the list of the 21 viral peptides with a combination of peptides from HBV, SARS CoV, Flu, CMV, HIV, and EBV.

**Supplementary table 2:**
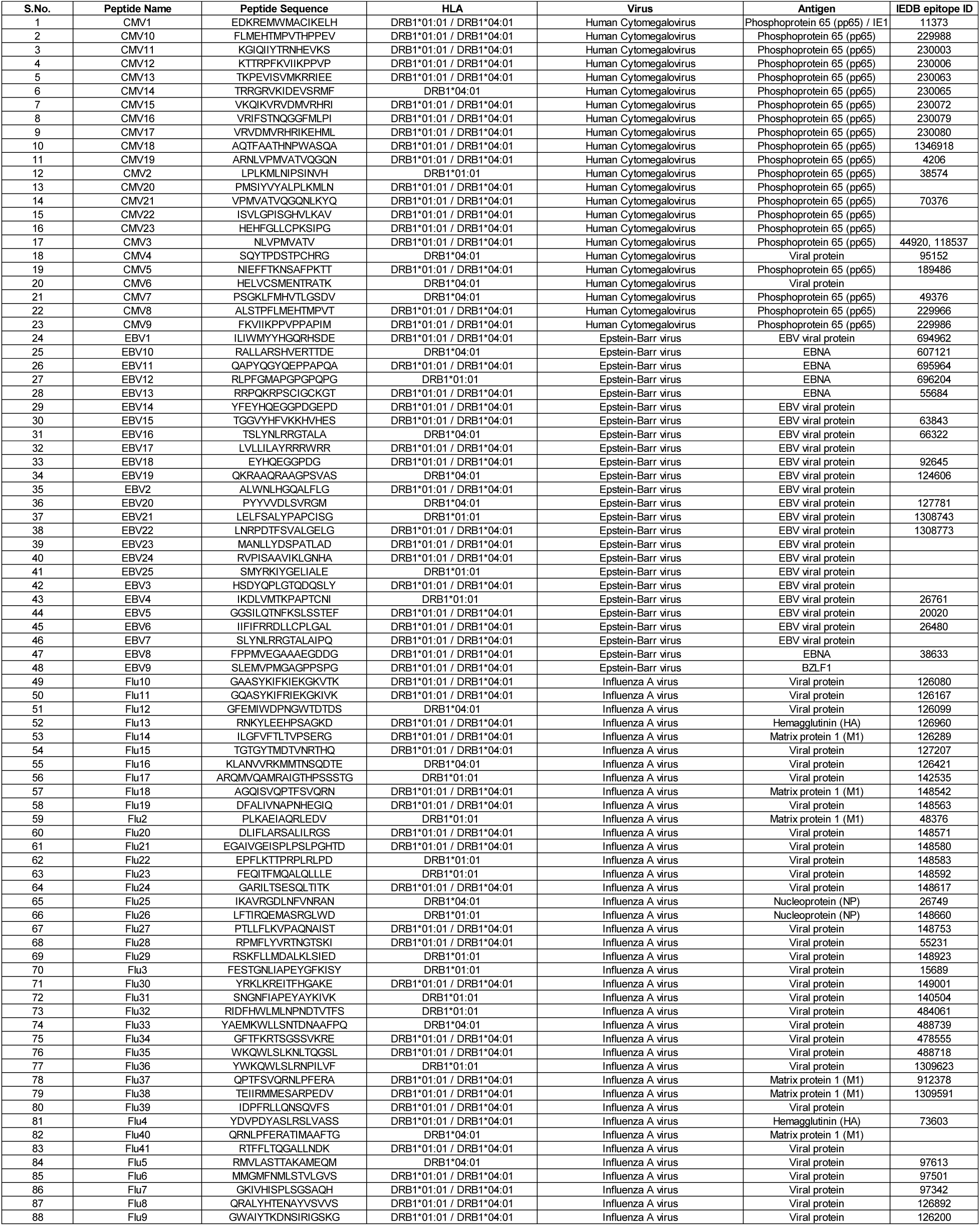
The table provides the list of the 88 expanded viral peptides for CMV, EBV, and Flu.

**Supplementary table 3:**
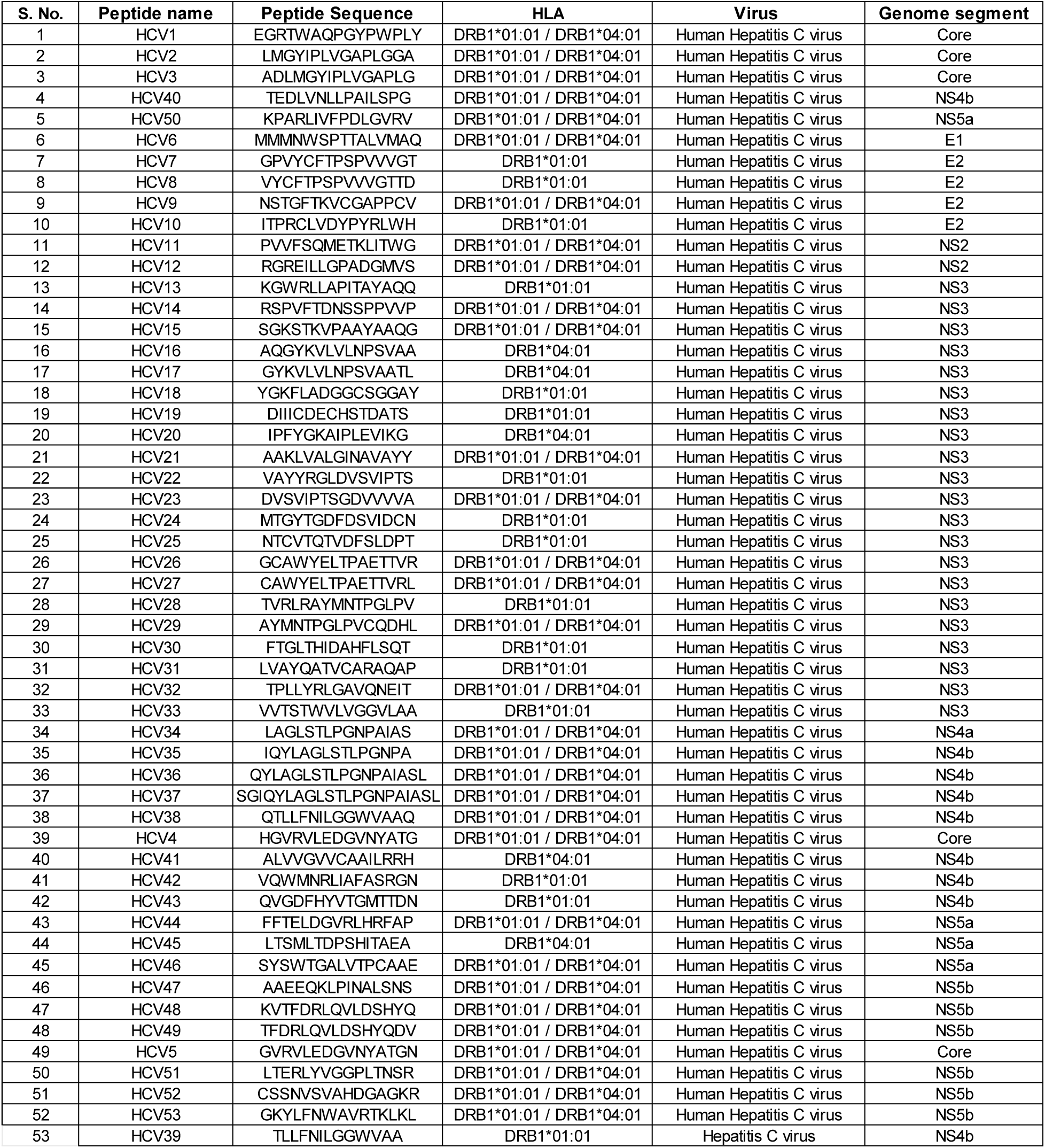
List of 53 HCV-specific peptides used for pMHCII multimer generation.

## Supplementary Figures

**Supplementary Figure 1:**
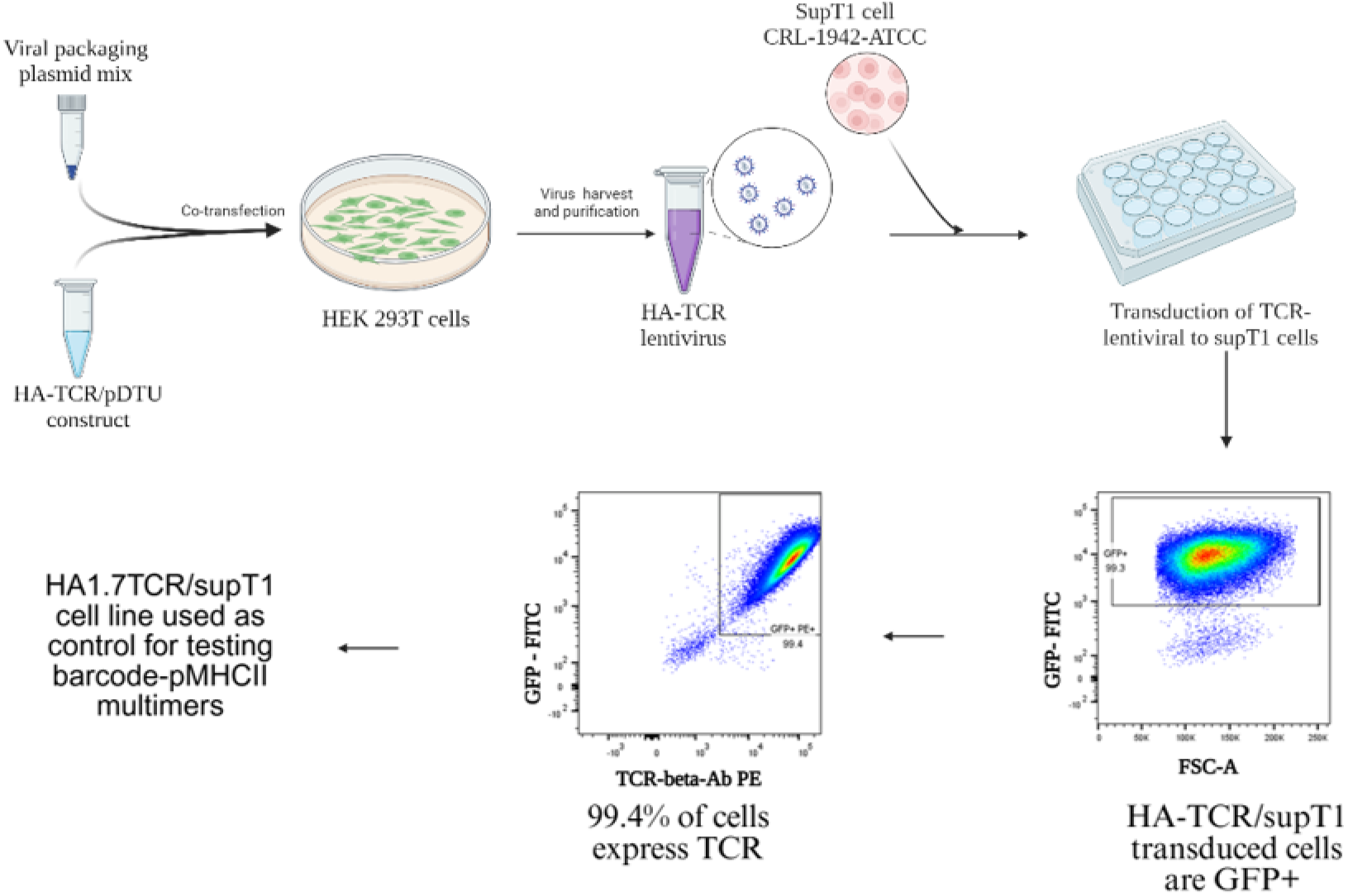
Engineering supT1 cells to express HA1.7 TCR against Flu1 (HA_306-318)_ epitope -. Engineered supT1 cells expressing HA1.7 TCR showed transduced cells as GFP^+^ with 99.3% transduction, and all transduced cells bind to the TCR-beta antibody shown as GFP^+^PE^+^.

**Supplementary Figure 2:**
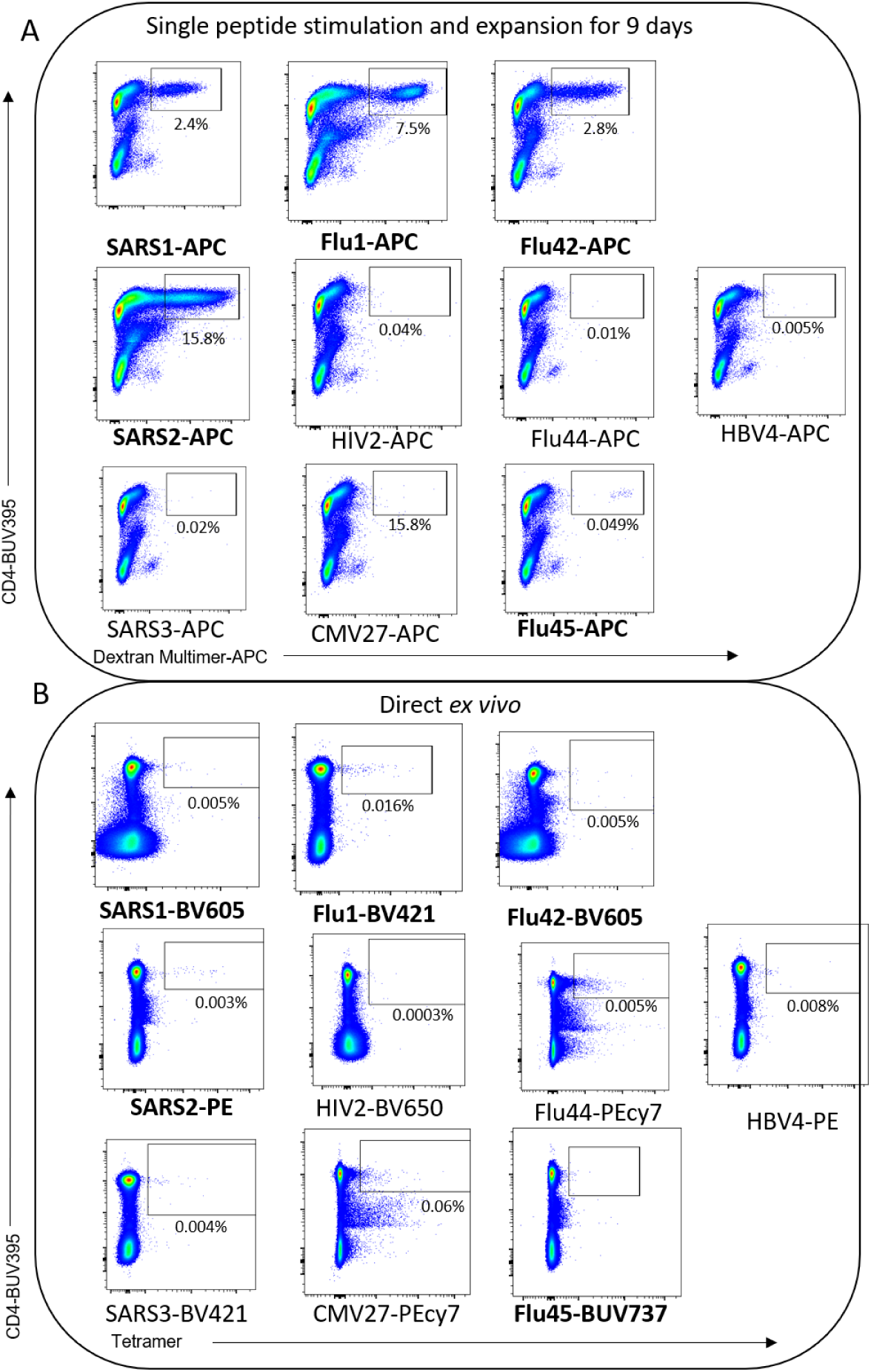
Multimer staining of PBMC HD1 as direct *ex vivo* and peptide expanded with individual peptides-. On top, flow cytometry plots showing the dextran multimer staining of different pMHCIIs of HD1 with single-peptide stimulated and expanded for 9 days. The peptide with which PBMC was expanded, was also respectively used to stain the cells as dextran multimers (mentioned below each flow plot). In the bottom, direct ex vivo screen of individual tetramers (peptide name and fluorophore marked in the bottom of each plot). Peptides where positive population was seen either in peptide-expanded or direct ex vivo PBMC were bolded.

**Supplementary Figure 3:**
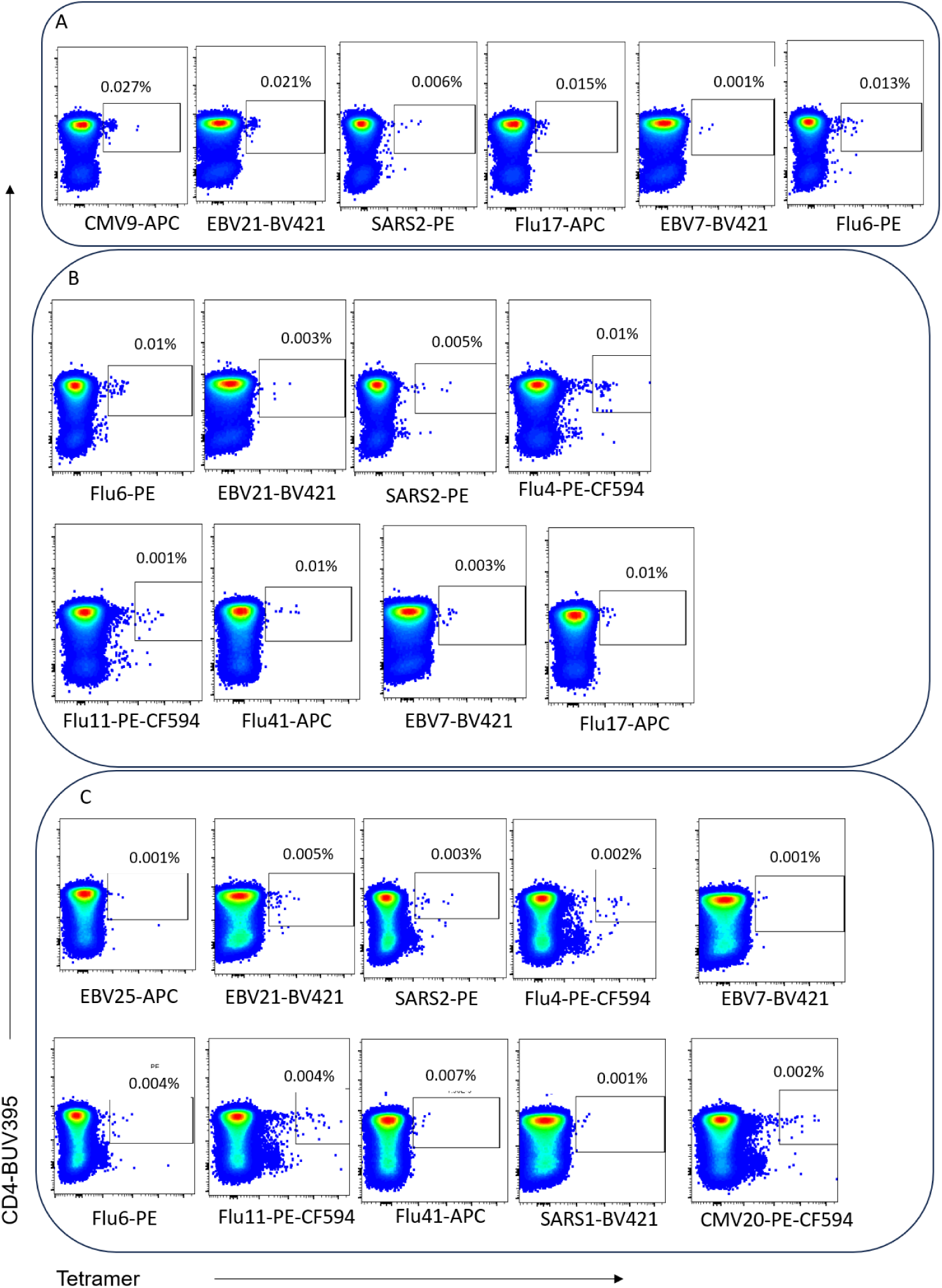
Validation of significant responses in healthy donors by conventional tetramer staining in direct *ex vivo* PBMC-. Flow cytometry plots showing the tetramer staining of different pMHCIIs on fluorophore-streptavidin (specified at the bottom for each plot) in direct *ex vivo* samples. A, B and C represents flow plots for donors HD6, HD9, and HD4 respectively.

**Supplementary Figure 4:**
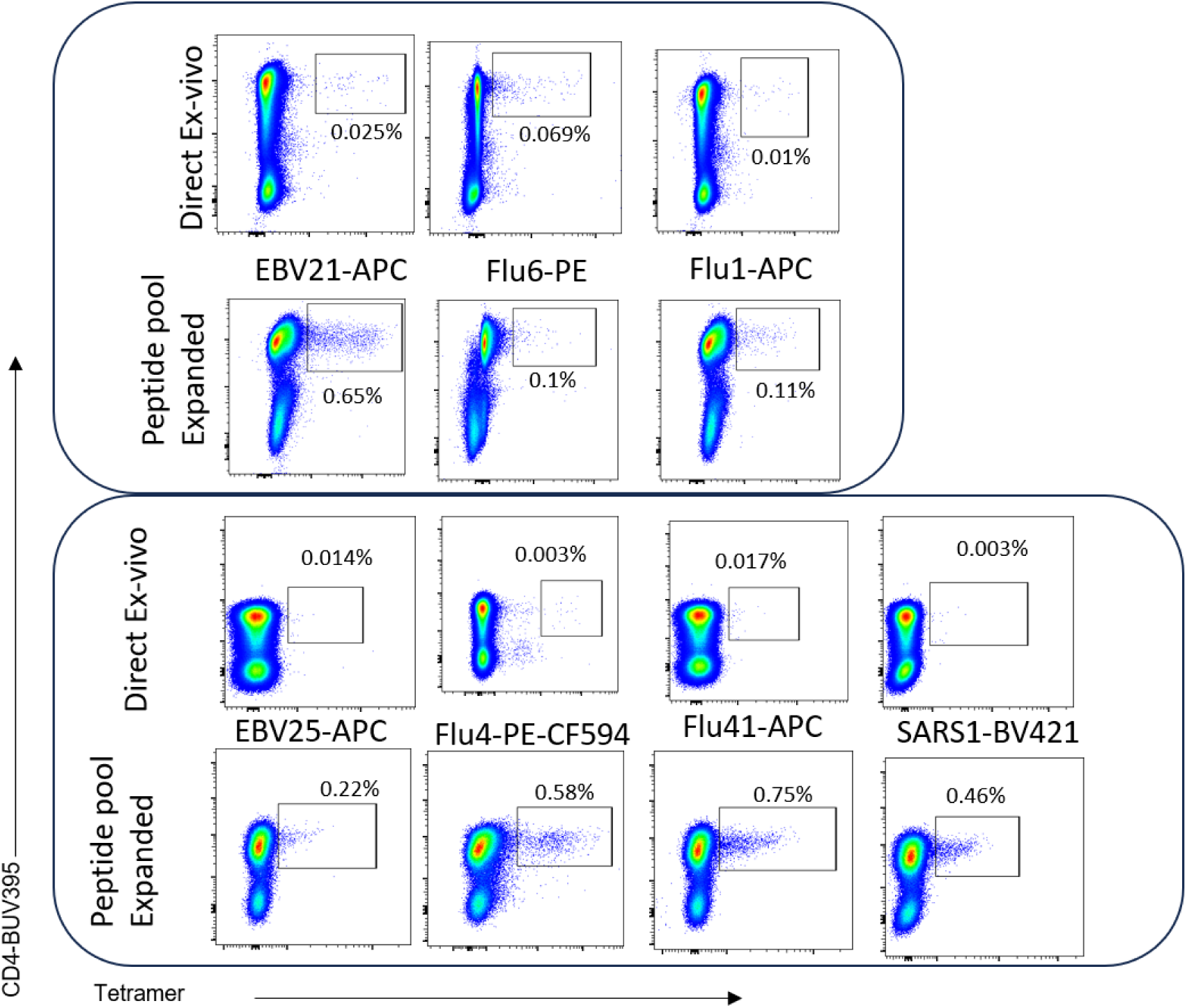
Validation of significant responses in HD2 by conventional tetramer staining. Flow cytometry plots showing the tetramer staining of different pMHCIIs on fluorophore-streptavidin (specified in the middle of direct ex vivo and peptide-pool expanded for each plot) in direct *ex vivo* and peptide-pool expanded samples of HD2.

**Supplementary figure 5:**
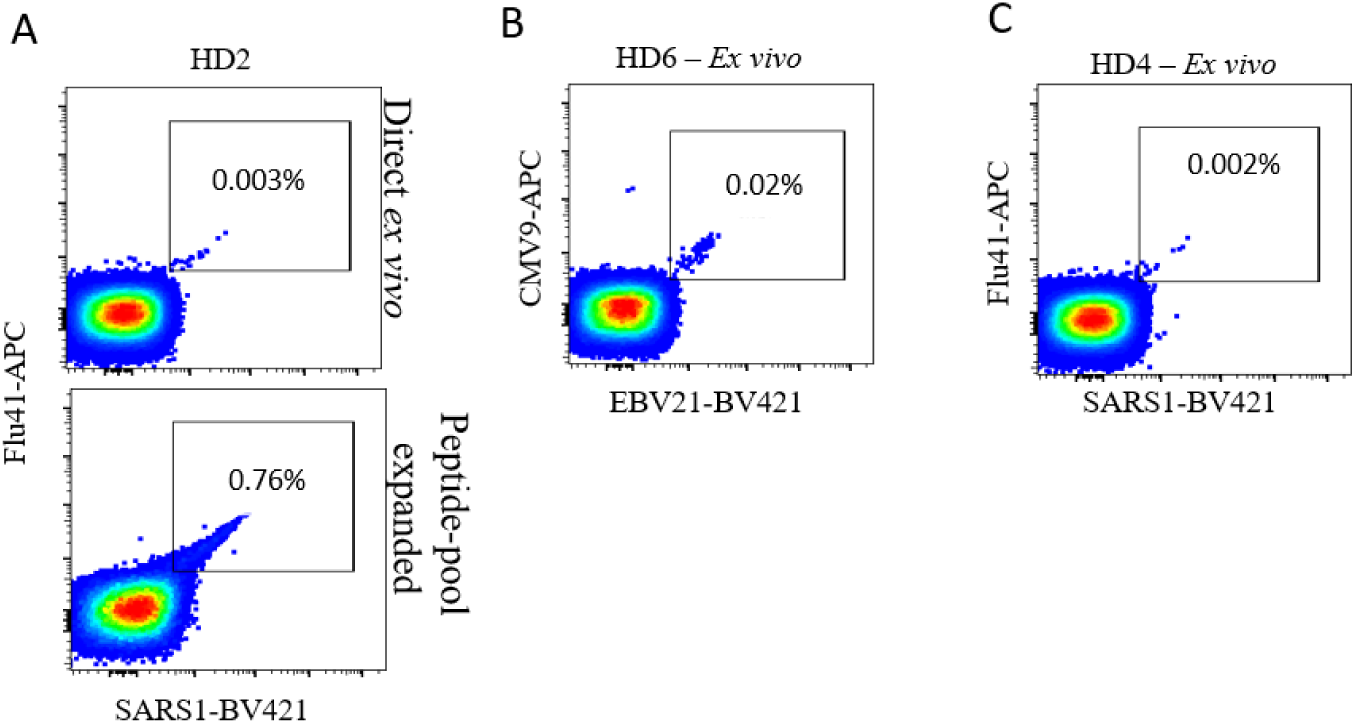
Cross reactive responses detected during tetramer validation – Flow plots representing cross reactivity between epitopes -. A and C shows cross reactivity between Flu41 and SARS1 pMHCII complexes in samples HD2 (direct ex vivo and peptide-pool expanded) and HD4 (direct ex vivo) respectively. B shows cross reactivity between CMV9 and EBV21 pMHCII in sample HD6.

**Supplementary figure 6:**
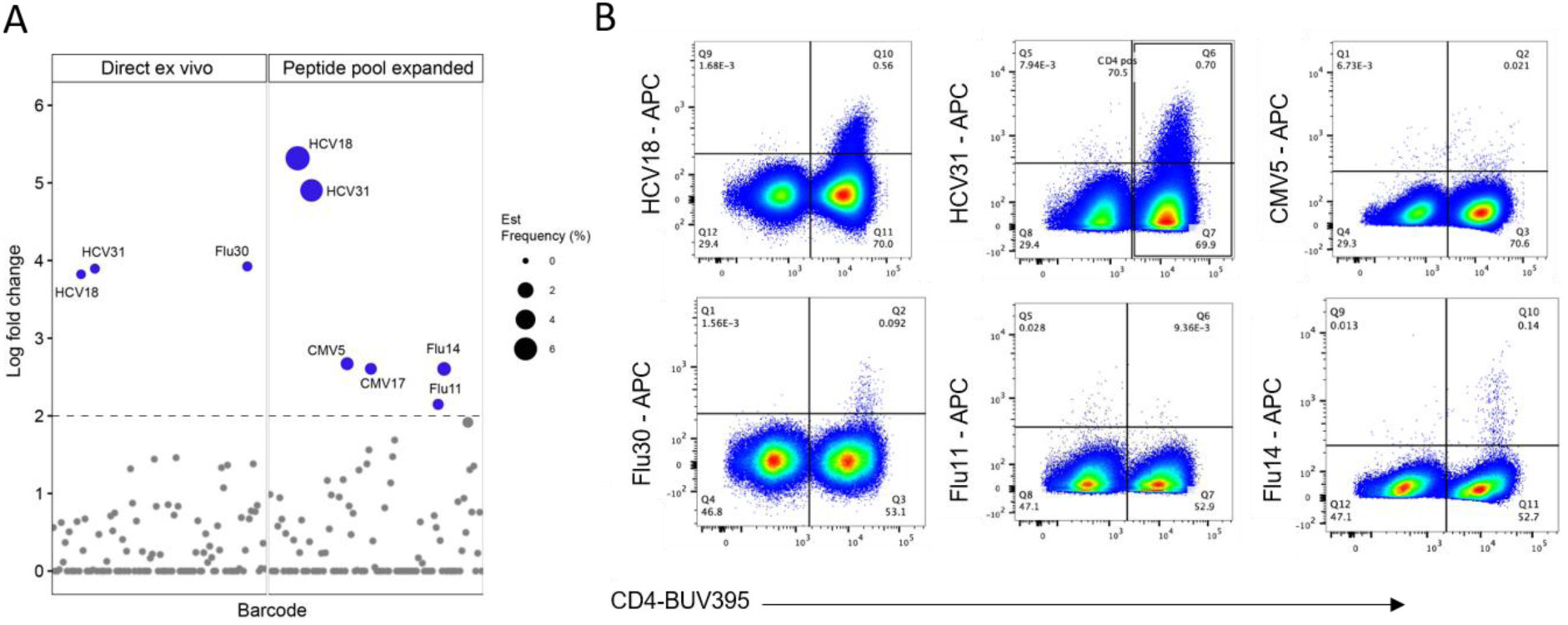
Screening of HCV participant 98-00A by barcoded-pMHCII multimers and validation of the responses-. A) Log fold change plot showing significant barcode-enrichment for the respective pMHCII multimers as both direct *ex vivo* and peptide-pool expanded. B) Conventional tetramer validation of the significant responses. The pMHCII-tetramer population is represented as %CD3 T cells in the top-right corner of each plots. In the flow plots representing HCV31-tetramer % CD4 T cell population specific to the pMHCII tetramer is shown in the top middle part of the plot.

**Supplementary figure 7:**
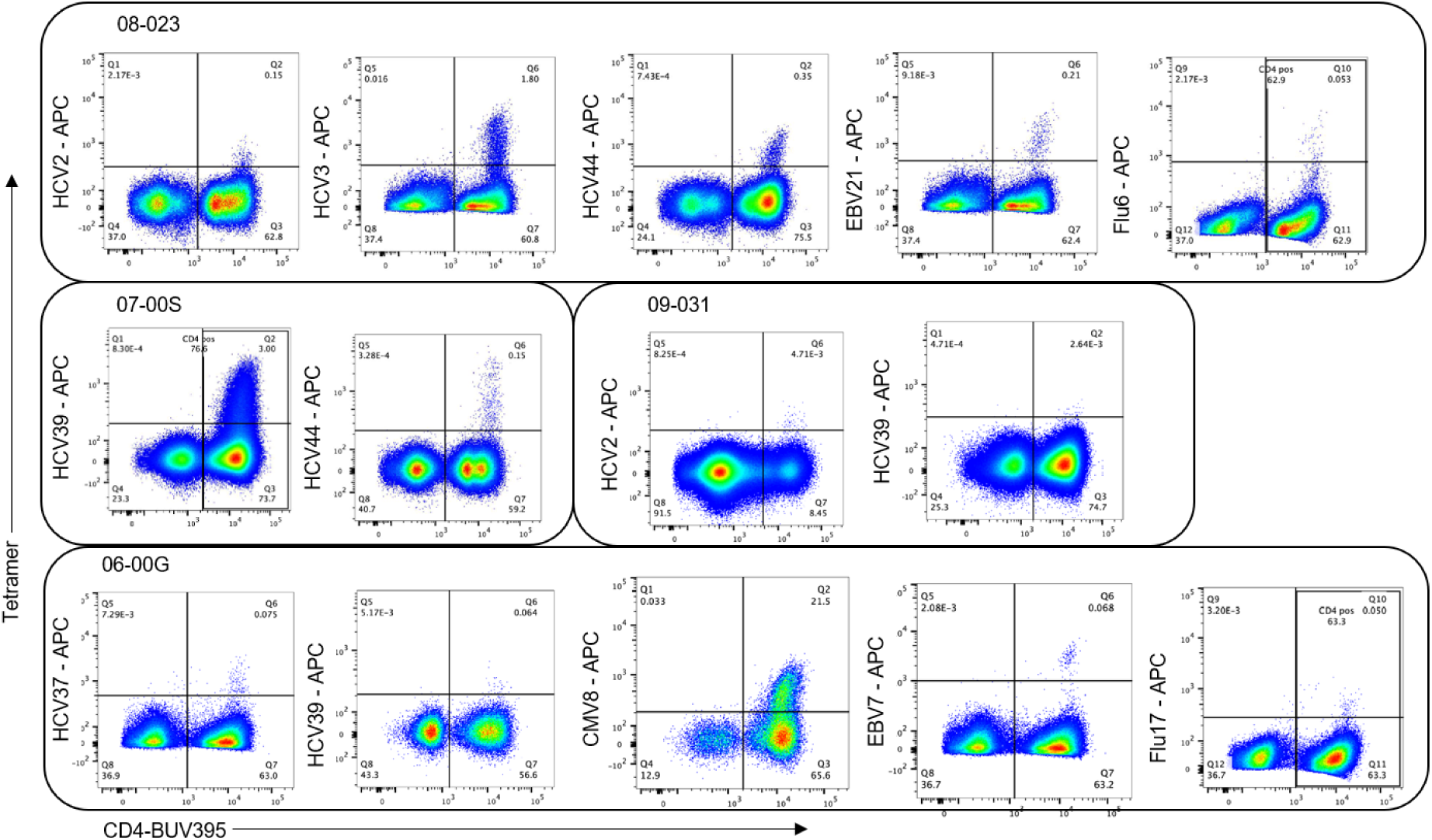
Tetramer validation of DRB1*01:01 restricted HCV participant-. Figure shows the flow plots of tetramer validation of HCV participants restricted to DRB1*01:01 grouped by the participant in boxes, with the participant ID written in the left top corner of the box. The pMHCII-tetramer population is represented as %CD3 T cells in the top-right corner of each plots. In the flow plots (HCV39 for participant 07-00S, Flu6 for participant 08-023, and Flu17 for participant 06-00G %CD4 T cell population specific to the pMHCII tetramer is shown in the top middle part of the plot.

**Supplementary figure 8:**
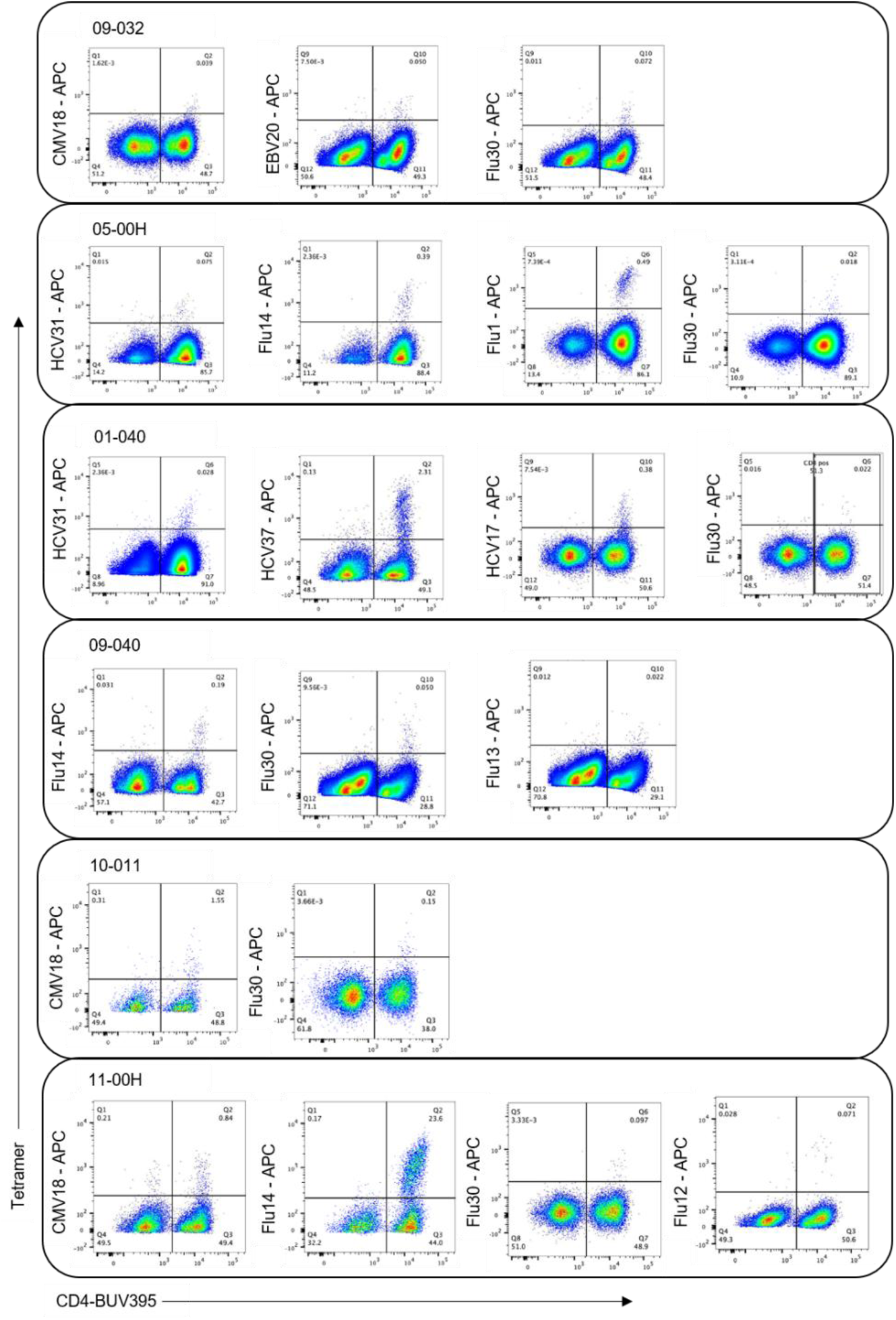
Tetramer validation of DRB1*04:01 restricted HCV participants-. Figure shows the flow plots of tetramer validation of HCV participants restricted to DRB1*04:01 grouped by the participants in boxes, with the participant ID written in the left top corner of the box. The pMHCII-tetramer population is represented as %CD3 T cells in the top-right corner of each plots. In the flow plots for Flu30 for participant 01-040, %CD4 T cell population specific to the pMHCII tetramer is shown in the top middle part of the plot.

**Supplementary figure 9:**
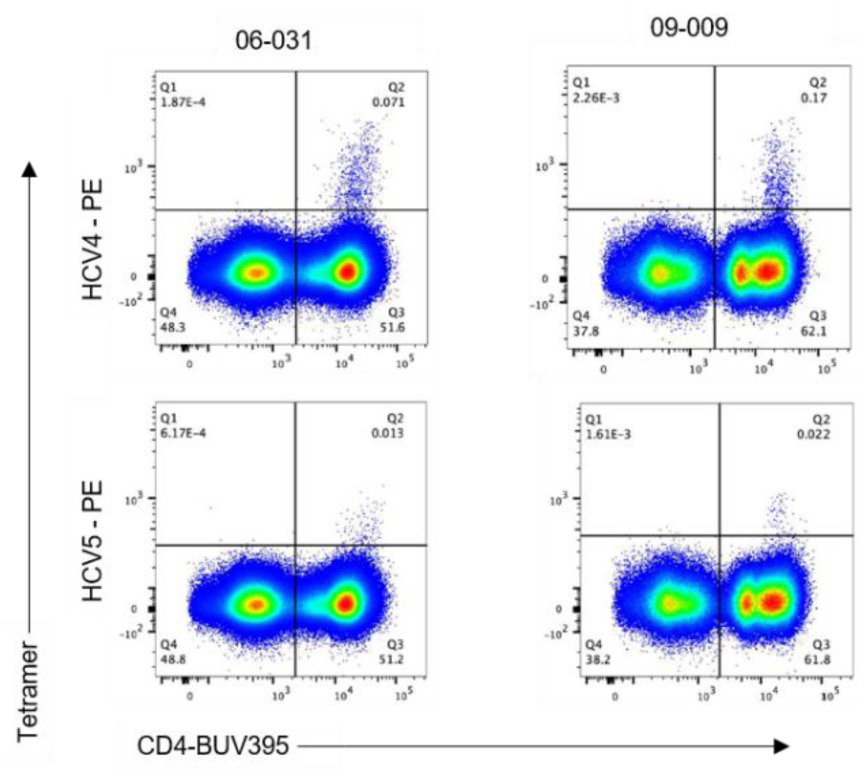
Tetramer validation of DRB1*04:01 restricted HCV4 and HCV5 peptides-. Figure shows the flow plots of tetramer validation HCV4 and HCV5 peptides in external HCV participants restricted to DRB1*04:01. The participant code is mentioned on top of the flow plot and the peptide-specific tetramer is mentioned on the y axis.

**Supplementary figure 10:**
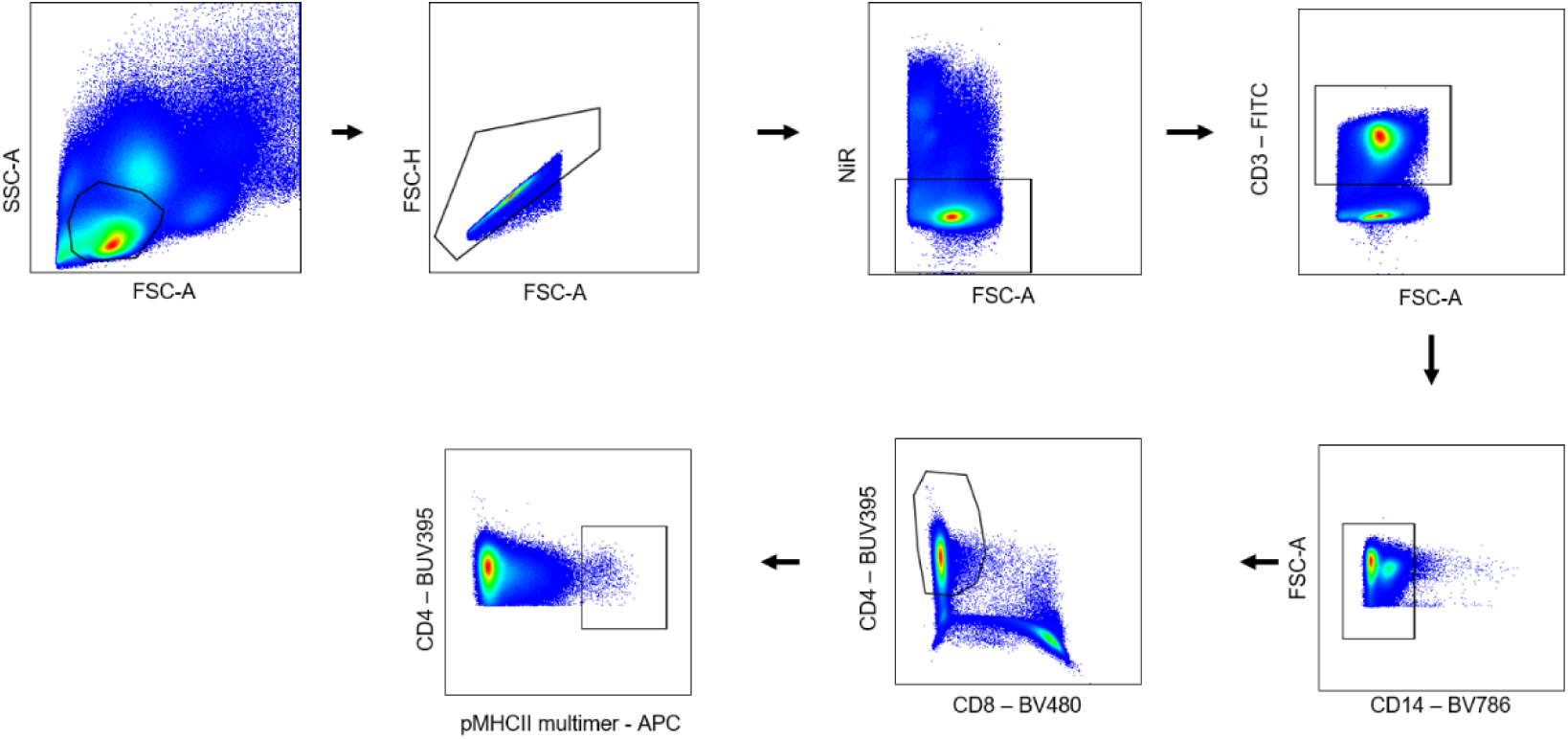
Sorting of barcoded-pMHCII multimer positive cells -. Flow cytometry gating strategy to sort multimer-positive CD4 T cells with a DNA-barcoded pMHCII multimer panel conjugated to APC.

**Supplementary figure 11:**
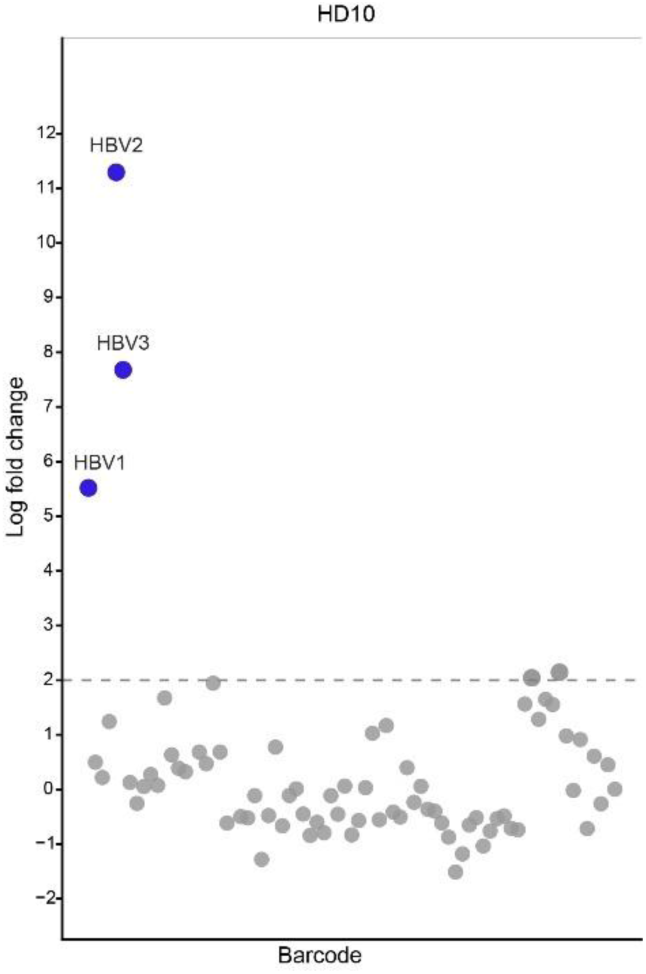
HLA-mismatch control in barcoded-pMHCII multimer experiment-. The plot displays the pMHCII-specific barcode enrichment in sample HD10 (HLA-mismatch control). The significant responses seen -HBV2, HBV3, and HBV1.

